# Scaffold modelling captures the structure-function-dynamics relationship in brain microcircuits

**DOI:** 10.1101/2021.07.30.454314

**Authors:** Robin De Schepper, Alice Geminiani, Stefano Masoli, Martina Francesca Rizza, Alberto Antonietti, Claudia Casellato, Egidio D’Angelo

## Abstract

Modelling brain networks with complex configuration and cellular properties requires a set of neuroinformatic tools and an organized staged workflow. We have therefore developed the Brain Scaffold Builder (BSB), a new modeling framework embedding multiple strategies for cell placement and connectivity and a flexible management of cellular and network mechanisms. With BSB, for the first time, the mouse cerebellar cortex was reconstructed and simulated at cellular resolution, using morphologically realistic multi-compartmental single-neuron models. Embedded connection rules allowed BSB to generate the cerebellar connectome, unifying a collection of scattered experimental data into a coherent construct. Naturalistic background and sensory-burst stimulation were used for functional validation against recordings *in vivo*, monitoring the impact of subcellular mechanisms on signal propagation and spatio-temporal processing and providing a new ground-truth about circuit organization for the prediction of neural dynamics.

## 2 Introduction

The relationship between structure, function and dynamics in brain circuits is still poorly understood posing a formidable challenge to neuroscience. The core issue is how to deal with the distribution and causality of neural processing across multiple spatio-temporal scales. In principle, a bottom-up model could take into account multi-modal datasets representing morphology, activity and connectivity of different cell populations and making it possible to build multi-scale realistic models, where microscopic phenomena propagate into large-scale network dynamics (Amunts et al., 2019; D’Angelo and Gandini Wheeler-Kingshott, 2017; Markram et al., 2015). If properly configured, these models can generate their own ground-truth by binding the many parameters, provided by independent measurements and intrinsically prone to experimental error, into a coherent construct. The bottom-up modelling strategy must be flexible enough to account for a large variety of neuronal features and network architectures, to incorporate new anatomical and physiological data when available, and to test various functional hypotheses (Brette et al., 2007). Moreover, the network must be scalable to the nature of the scientific question and to efficiently interface with available simulation platforms, e.g. NEURON (Hines and Carnevale, 1997) and NEST (Gewaltig and Diesmann, 2007). There are modelling frameworks with solutions to most common problems (Dai et al., 2020; Dura-Bernal et al., 2019; Gratiy et al., 2018) but there is still a need for toolkits taking into account the location, morphologies and rotation of cells and using these properties to determine connectivity. Here, we introduce the Brain Scaffold Builder (BSB), an advanced framework for neuronal circuit modelling, with specific modules for network reconstruction and simulation. The “scaffold” design allows an easy model reconfiguration reflecting variants across brain regions, animal species and physio-pathological conditions without dismounting the basic network structure.

We have taken the moves from the cerebellar cortical microcircuit (D’Angelo et al., 2016) that has inspired foundational theories on brain functioning (Marr, 1969) but challenges realistic computational modelling. The main limitation of previous models was that connectivity was independent from neuronal morphology (Casali et al., 2019, 2020; Solinas et al., 2010) preventing a direct link between microcircuit structure, function and dynamics. With BSB, we have generated the first computational model of the cerebellar microcircuit including a layered volume with variegate neuron types characterized by different density, morphology and orientation, which were wired through a connectome defined by the anisotropy of dendritic and axonal processes through principled rules. The insertion of single cell multicompartmental models and dynamic synapses (Masoli et al., 2020a, 2020b; Masoli and D’Angelo, 2017; Rizza et al., 2021) was key to simulate network dynamics in response to naturalistic inputs (Ramakrishnan et al., 2016; Rancz et al., 2007; Roggeri et al., 2008). This work provides a new structural and functional ground-truth for the cerebellar circuit and implies the effectiveness of the BSB multi-scale modelling strategy to different brain areas.

## 3 Results

### 3.1 The Brain Scaffold Builder (BSB)

BSB is a modelling framework designed to study neural structure and computation under realistic biological constraints and operates through a sequence of independent steps: network description, reconstruction and simulation (Fig. 1a). The network volume is first defined along with the cell types, then BSB proceeds through cell placement and connectivity reconstructing the microcircuit network (Fig. 1b, c). Finally, network activity is simulated and the results visualized. BSB allows to choose among pre-designed algorithms and to plug-in user-defined ones. BSB allows to replace neurons and synapses and modify their parameters (Fig. S1). BSB includes interfaces for different simulators (e.g. Neuron and NEST) and can be installed on any devices where Python is available (*python -m pip install bsb*), is open source with packaged and installable distribution, source code documentation and topical tutorials (Fig. S1). In aggregate, BSB is conceived as an efficient modelling tool for brain microcircuit modeling and simulation.

**Figure 1.**
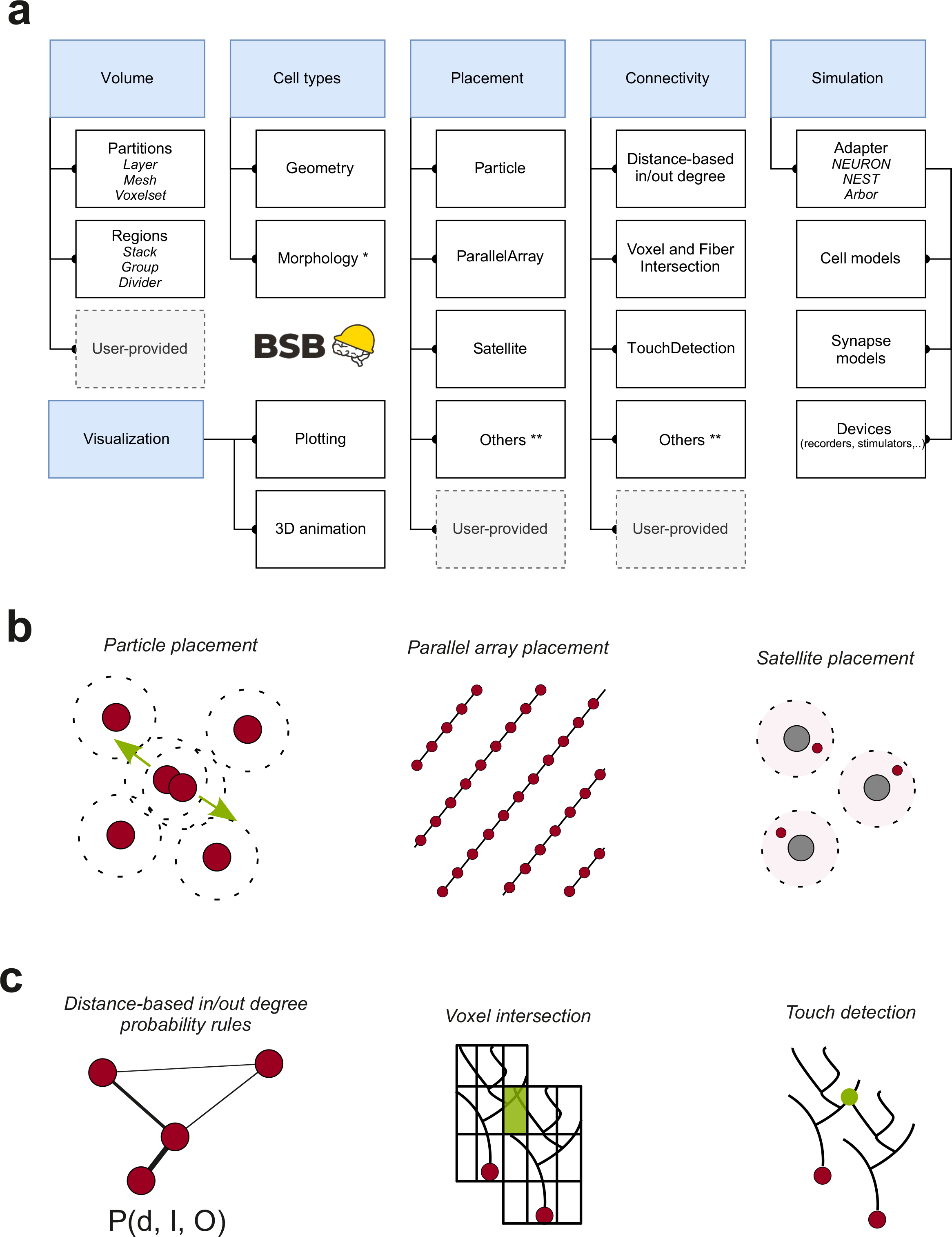
The Brain Scaffold Builder. **(a)** Core BSB operations. In the reconstruction phase, BSB proceeds by sequentially defining the network *volume*, *cell types*, *cell placement, cell* c*onnectivity*. Once neurons are positioned, their geometries/morphologies are imported and connection rules allow to wire them up and to build the network connectome. In the simulation phase, *neuron* and *synapse models* are linked to simulators, like NEURON and NEST, by specific adapters and interfaced to a set of *devices* for stimulation and recording. In the post-simulation phase, graphic tools are made available for data representation. This workflow is applicable to any kind of brain neuronal network. **(b)** Infographic representations of the main *placement strategies* available in BSB, using kd-tree partitioning of the 3D space (particle placement, parallel array placement, satellite placement). **(c)** Infographic representations of the main *connection strategies* available in BSB: distance-based in/out degree probability functions, voxel (or fiber) intersection based on voxelization of morphologies, touch detection. It should be noted that BSB embeds strategies also for the case in which morphologies are not available and that user-defined routines can be easily added. The application of BSB to the case of the cerebellar network microcircuit is illustrated in the next figures.

### 3.2 Cerebellar network reconstruction

BSB was applied first to the mouse cerebellar cortical network, which has a geometrically organized architecture that has been suggested since early to imply its computational properties (D’Angelo et al., 2016; Marr, 1969). The reconstruction and simulation of a network volume of 17.7 10^-3^ mm^3^ is reported, including the following cell and fiber types: mossy fiber (*mf*), glomerulus (Glom), granule cell (GrC) with ascending axon (*aa*) and parallel fiber (*pf*), Golgi cell (GoC), Purkinje cell (PC), molecular layer interneurons (MLI) comprising stellate cell (SC) and basket cell (BC).

#### 3.2.1 Neuron placement

The network elements summed up to 29’230 neurons (GrC, GoC, PC, SC, BC) plus 2’453 other elements (*mf*, Glom), which were placed in the network volume according to anatomical data (references from (Casali et al., 2019; D’Angelo et al., 2016; Sultan and Bower, 1998)) through procedures of *particle placement* and *parallel array placement* (Fig. 2a). The density values matched the targets given in the configuration file, nearest neighbor distances always exceeded cell diameter, and radial density functions demonstrated the homogeneity of cell distribution (Fig. S2). The algorithms produced realistic cell positioning avoiding unphysiological lattice-like structures (Töpperwien et al., 2018).

**Figure 2.**
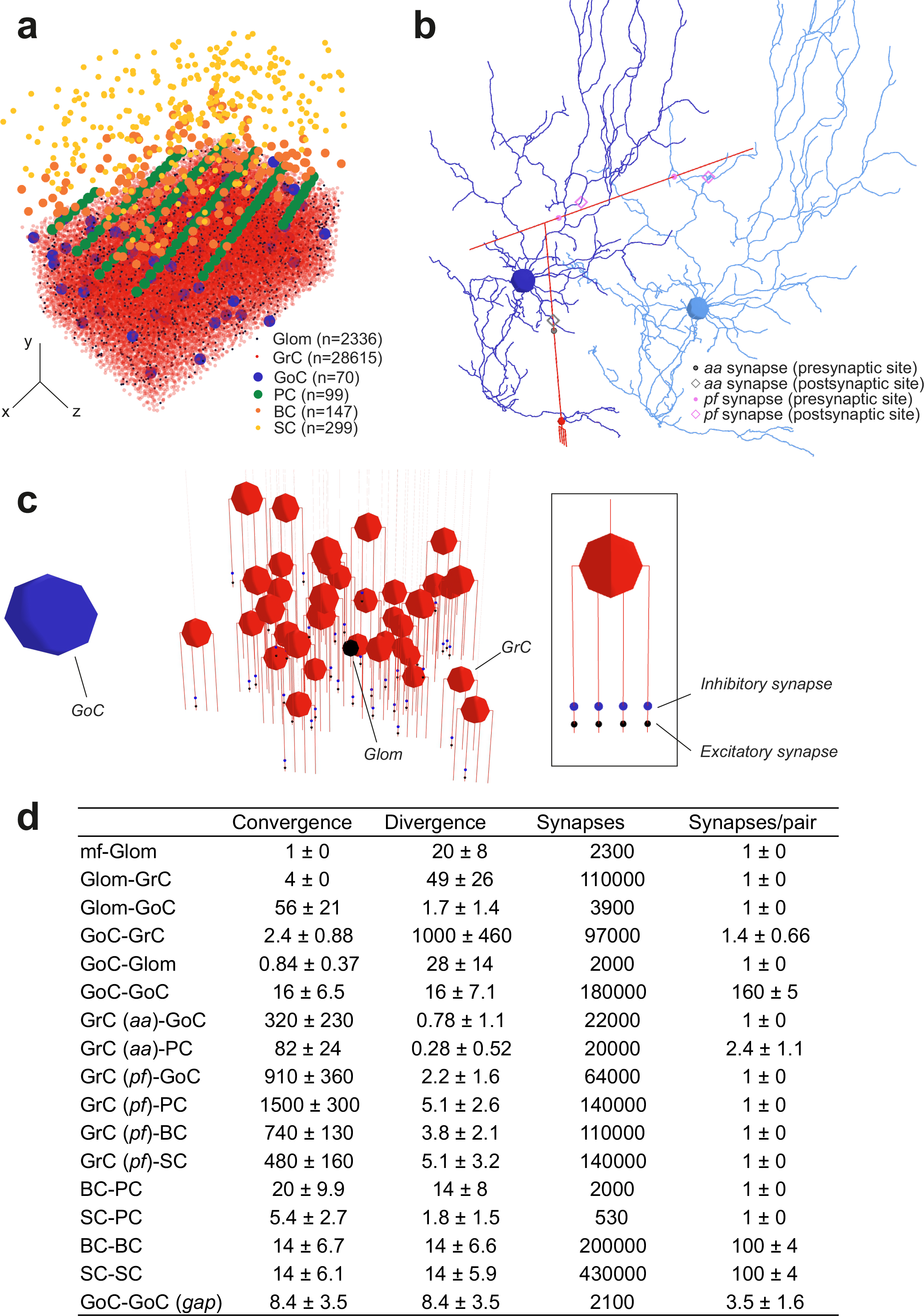
Reconstruction of the microcircuit of cerebellar cortex. **(a)** Positioning of cell bodies in a 3D slab (300 x 295 x 200 μm^3^) of mouse cerebellar cortex. Cell numbers are indicated (the symbols reflect soma size). In this and the following figures, the *xyz* reference system is defined by *x-y* (sagittal plane), *x-z* (axial plane), *z-y* (coronal plane), as in standard anatomical representations. Thus, *y* measures cortex thickness (*aa* direction), while *z* identifies the major lamellar axis (*pf* direction). **(b)** Example of 3D morphologies illustrating GrC-GoC connections through *aa* and *pf*. Two GoCs and one GrC are shown: the synapse along *aa* is identified by *touch detection*, while synapses along *pf* are identified by *fiber intersection*. **(c)** Glom-GrC and GoC-GrC connections. A Glom contacts a group of 38 GrCs forming an excitatory synapse on the terminal compartment of 1 of their 4 dendrites. The Glom, in turn, is contacted by a GoC nearby, which forms inhibitory synapse on the pre-terminal dendritic compartment of the same GrCs. The inset shows a GrC with 1 excitatory synapse and 1 inhibitory synapse on each dendrite. **(d)** The cerebellar cortical connectome generated by BSB reporting convergence (on the postsynaptic element), divergence (from the presynaptic element), total number of synapses, and number of synapses for each connected pair.

#### 3.2.2 Neuron connectivity

The network connections summed up to 1’500’000 chemical synapses and 2’100 electrical synapses. The connectivity procedure combined probabilistic and geometric rules that were chosen depending on available data and network architecture (Fig. 2b,c) and yielded the metrics of cerebellar the connectome (Fig. 2d).

The connectivity of *mf* and Glom was accounted for by *distance-based in/out degree* algorithms. BSB generated local anisotropic Glom clusters extending 60 µm along the parasagittal and 20 µm along and transverse plane (Sultan, 2001), yielding ∼20 Gloms per *mf* (Billings et al., 2014). Imposing that each GrC sends its 4 dendrites to Gloms belonging to different *mfs* within about 30 µm, BSB yielded 49 GrCs per Glom on average (Houston et al., 2017; Jakab and Hámori, 1988) (Fig. 2b).

The connectivity of GoCs posed more complex issues. (i) GoCs receive excitatory inputs from *mf*s (Tabuchi et al., 2019). In BSB, each GoC received excitation from 56 different Gloms and each Glom collected basolateral dendrites from ∼2 GoCs. (ii) Given that GoC-GrC synapses are inside the glomeruli, each GrC dendrite receives inhibition from a GoC whose axon reaches the Glom contacting that dendrite (Hamori et al., 1997; Mapelli et al., 2014). In BSB, each GrC had 4 inhibitory synapses, one per dendrite, mostly originating from different GoCs (Fig. 2c). (iii) GoCs were reported to receive ∼400 *aa* synapses on basolateral dendrites and ∼1200 *pf* synapses on apical dendrites (Cesana et al., 2013). In BSB, *touch detection* identified 320 *aa* synapses and *fiber intersection* identified 910 *pf* synapses per GoC, all from different GrCs (Fig. 2c). (iv) GoCs make GABAergic synapses onto other GoCs (Hull and Regehr, 2012). In BSB, *voxel intersection* identified inhibition from other 16 GoCs on the basolateral dendrites and subsequent functional calibration implied ∼160 synapses per pair (see below). (v) Finally, there are 2-4 gap-junctions per GoC pair (Szoboszlay et al., 2016). *Voxel intersection* identified 8.4 GoCs that form gap junctions on other GoCs, with ∼3.5 gap-junctions per pair.

The number of *pf -*PC synapses may range up to ∼100’000 (Hoxha et al., 2016), many of which would be silent (Isope and Barbour, 2002). Based on spines density (Parajuli and Koike, 2021) and the total length of PC dendritic tree (Masoli and D’Angelo, 2017), the number of possible *pf* synapses was estimated to be 15’000-20’000, with a minimum of ∼100 synapses needed to generate a simple spike (Walter and Khodakhah, 2006). Here, limited to our 200-μm *pf* length, *fiber intersection* identified 1’500 *pf* synapses per PC. T*ouch detection* identified 82 *aa* synapses per PC with ∼2.4 *aa* synapses / PC.

The number of synapses formed by *pf*s on MLIs is not well defined. *Fiber intersection* identified 480 *pf* synapses per SC and 740 per BC. Moreover, MLIs form connections with other MLIs of the same type (Kondo and Marty, 1998). *Voxel intersection* identified 14 SC-SC and 14 BC-BC connections, forming ∼100 synapses per pair. In turn, the collaterals of a SC axon, mainly extending on the coronal plane, innervate multiple PCs (Ango et al., 2008). V*oxel intersection* identified 5.4 SCs contacting each PC (Fig. S3). Moreover, the BC axons typically extend in the sagittal plane and innervate 7-10 PCs (Ango et al., 2008), ending up with 3-50 baskets around the PC soma (Blot and Barbour, 2014). *Voxel intersection* identified ∼20 BC synapses per PC and ∼14 PCs per BC. Parameter predictions were further assessed and tuned through functional simulations (see below).

### 3.3 Cerebellar network simulations

Network simulations were carried out using detailed neuronal and synaptic models written in NEURON for GrC (Masoli et al., 2020b), GoC (Masoli et al., 2020a), PC, (Masoli and D’Angelo, 2017), SC and BC (Rizza et al., 2021). Local microcircuit responses to input patterns were tracked back to individual neurons and used to follow signal propagation with unprecedented resolution. All simulations were carried out in the presence of background noise to improve comparison with recordings *in vivo*. The emerging spatio-temporal dynamics provided functional model validation beyond constructive validity based on internal connectivity and single neuron responses (Movie S1).

#### 3.3.1 Resting state activity of the cerebellar network

A random input at low frequency (4 Hz Poisson) on all *mf*s (Rancz et al., 2007) was used to simulate the cerebellar network in resting state *in vivo*. Since anatomical data about the connectivity of cerebellar neurons are incomplete, but their resting discharge frequency is known, we finetuned the number of connections against target parameter values of resting discharge in GrCs (Wilms and Häusser, 2015), GoCs, (Forti et al., 2006; Masoli et al., 2020a; Solinas et al., 2007), PCs (Arancillo et al., 2015), SCs and BCs (Barmack and Yakhnitsa, 2008; Jirenhed et al., 2013; Kim and Augustine, 2020).The turning point was to calibrate GoC-GoC inhibition, which influenced resting-state activity of the entire network. Since the synaptic conductance (∼ 3200 pS) and the number of interconnected GoCs (about 15) are known (Hull and Regehr, 2012), we tuned the number of GoC-GoC synapses until basal discharge frequency was normalized. Eventually, the background frequency of all cerebellar neuron types fell in the ranges reported *in vivo* in anaesthetized rodents (*mf*s: 4.2 ± 2.6 Hz; GrCs: 0.81 ± 1.3 Hz; GoCs: 19 ± 15 Hz; PCs: 31 ± 1.6 Hz; BCs: 11 ± 5.1 Hz; SCs: 9.4 ± 12 Hz).

Background *mf* activity is known to generate synchronous low-frequency oscillations in the granular layer (Hartmann and Bower, 1998). Indeed, in the model, the FFT of GoC and GrC firing revealed a synchronous oscillatory behaviour in the theta band, with the first harmonic peaking at 9.7 Hz (Fig. 3a). When gap junctions were switched off, the regularity of the oscillation decreased and the first FFT harmonic moved out of theta band (Dugué et al., 2009).

**Figure 3.**
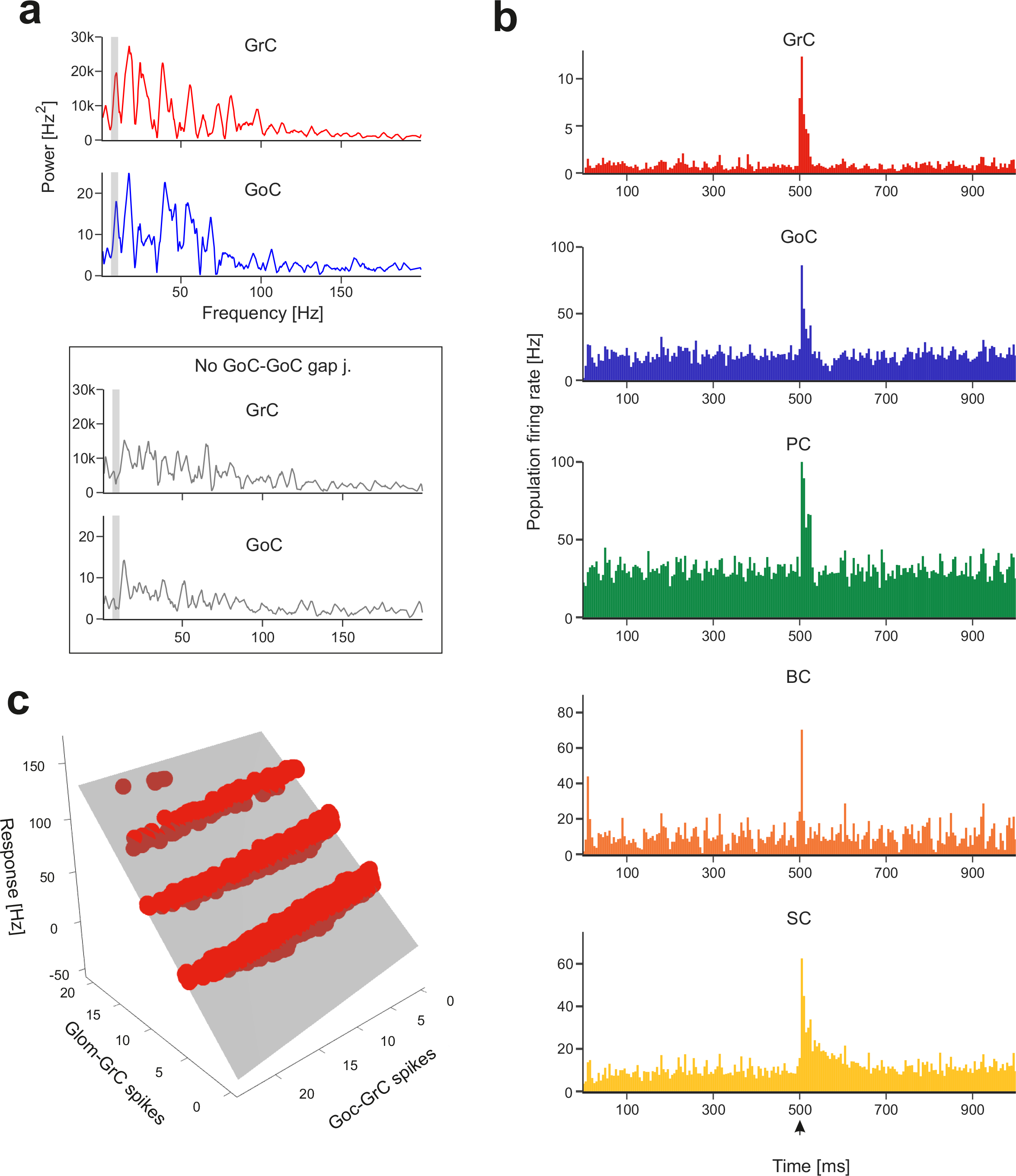
Network responses to background noise and *mf* bursts. **(a)** Power spectra of GrC and GoC activity were computed with Fast Fourier Transform (FFT) of spike time series (total population spike-counts in 2.5 ms time-bins). The periodicity of peaks in power spectra reveals synchronous low-frequency oscillations in the granular layer. The grey curves represent the power spectra when GoC-GoC gap junctions were disabled, showing a marked decrease in periodicity. The grey bands correspond to mouse theta-band (5-10 Hz). **(b)** The Peri-Stimulus-Time-Histograms (PSTH) of each neuronal population show the effect of a localized *mf* burst (arrowhead) emerging over background noise. The PSTHs show number of spikes/5 ms time-bins normalized by the number of cells (average of 10 simulations). **(c)** Example of multiple linear regression of GrC responses (firing rate during 40 ms after stimulus onset) against the number of synaptic spikes from Gloms and GoCs (the grey surface is the fitted plane to the points).

#### 3.3.2 Impulsive response of the cerebellar network

Short stimulus bursts were delivered to a bundle of 4 *mf*s connected to ∼80 Gloms to emulate whisker/facial sensory stimulation *in vivo* (Rancz et al., 2007; Wilms and Häusser, 2015). The bursts propagated through the network temporarily raising neuronal firing (Fig. 3b). The relationship between the number of spikes at afferent synapses and the response frequency to the *mf* burst was robustly captured by multiple linear regression (Fig. 3c; Fig. S4a; Fig. S5).

##### 3.3.2.1 GrC responses

Fundamental predictions on how GrCs respond to incoming bursts derive from current clamp recordings *in situ* (D’Angelo et al., 1995) and simulations (Masoli et al., 2020b), which revealed the role of synaptic receptors and ionic channels. In BSB simulations, bursts on a collimated *mf* bundle activated a dense cluster of GrCs (Casali et al., 2020; Diwakar et al., 2011; Roggeri et al., 2008) (Movie S1). The relationship between the number of input spikes (both at GoC-GrC and Glom-GrC synapses) and GrC response frequency unveiled 4 groups of GrCs with a corresponding number of synaptically activated dendrites (Fig. 3c). The number of GrC spikes, first spike latency and dendritic [Ca^2+^]_in_ correlated with the number of active dendrites (NMI = 0.71, 0.86, 0.59, respectively) (Fig. 4b).

**Figure 4.**
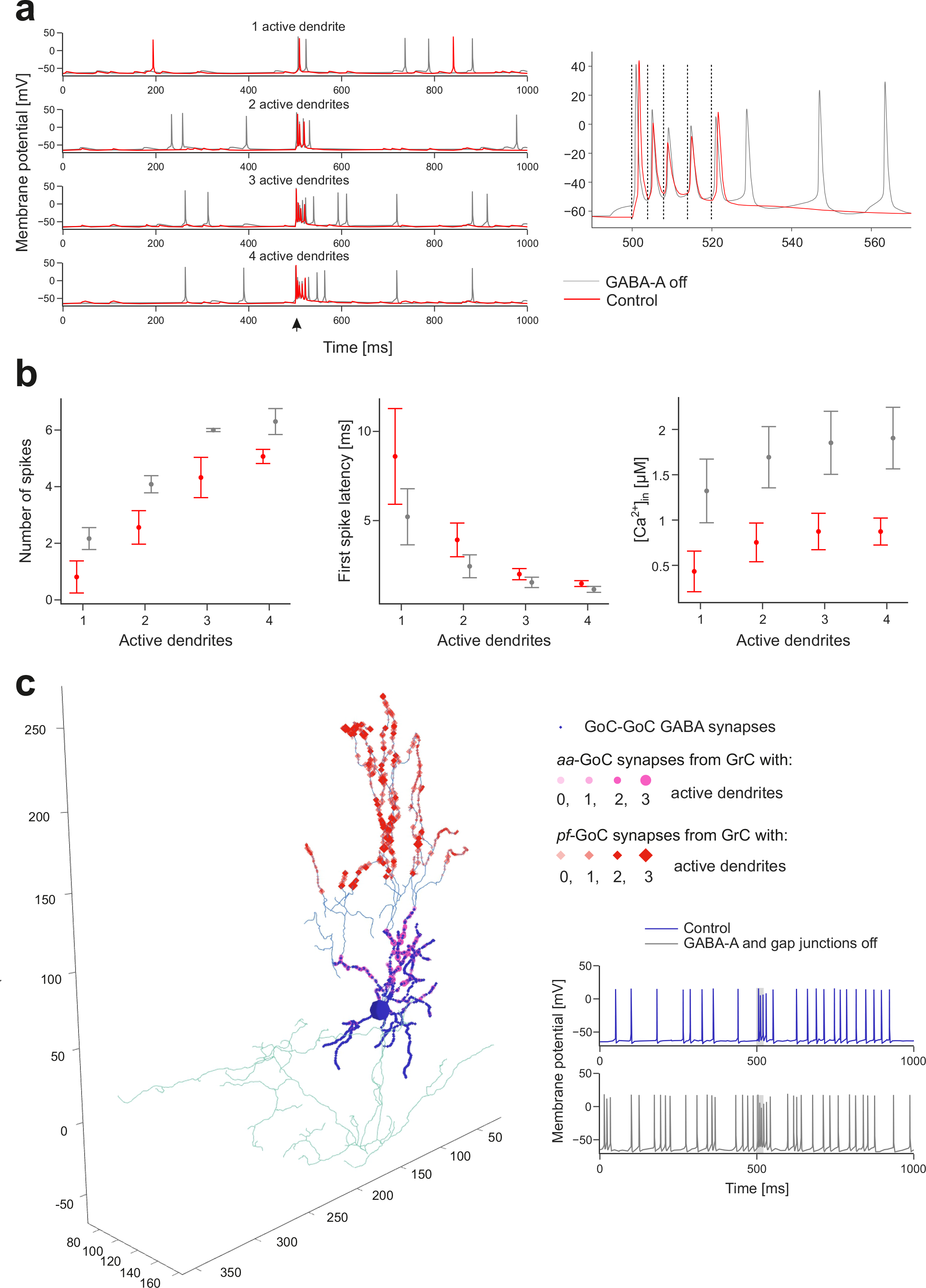
Granular layer activation. **(a)** Membrane potential of 4 representative GrCs with 1 to 4 dendrites activated by the *mf* burst (20ms@200Hz over background noise, onset indicated by arrowhead) in control condition and after GABA-A receptors blockade. The burst response is enlarged on the right to highlight spike-timing (dashed lines indicate *mf* spikes). **(b)** Number of spikes (measured in the 40 ms from *mf* burst onset), first spike latency, and dendritic [Ca^2+^]_in_ (measured in the 500 ms from *mf* burst onset) in subgroups of GrCs with the same number of activated dendrites (mean±sd). The graphs compare responses in control and during GABA-A receptors switch-off. **(c)** Synapses of a GoC activated by GrCs. The GABAergic synapses from other GoCs are on basal dendrites, *aa* synapses are on basal dendrites, *pf* synapses are on apical dendrites. In this example, the GoC receives 30% of its *aa* synapses and 6% of its *pf* synapses from GrCs with at least 2 active dendrites. Traces on the right show the GoC membrane potential in response to a *mf* burst (same stimulation as in (a), grey band) in control and during GABA-A receptors and gap junctions switch-off.

When the inhibitory mechanisms (comprising transient and persistent inhibition) were switched off to simulate a pharmacological GABA-A receptor blockade, GrC (i) baseline frequency increased, (ii) a tail discharge appeared after the burst, (iii) responses including more spikes appeared, (iv) the first spike latency decreased, and (v) response variability decreased (Fig. 4a,b), but the number of GrC spikes, first spike latency and dendritic [Ca^2+^]_in_ still correlated with the number of active dendrites (NMI = 0.79, 0.85, 0.61, respectively) (Fig. 4b). Interestingly, inhibition caused a reduction in the number of active GrCs (e.g. those firing >= 1 spike in the first 40 ms were 3390 ± 431 before and 8348 ± 1724 with GABA-A switch-off; n=10 simulations; p < 0.001, unpaired *t*-test) but enriched the spike pattern, as predicted theoretically (Eccles et al., 1967; Marr, 1969).

Recordings *in vivo* disclosed precise integration of quanta and high-fidelity transmission (Arenz et al., 2008; Chadderton et al., 2004; Ishikawa et al., 2015; Powell et al., 2015; Rancz et al., 2007). In BSB simulations, GrCs receiving maximum excitation generated one action potential for each spike of the input burst, with short latency (< 2 ms), and thus faithfully followed the input up to 250 Hz (Fig. 4a).

##### 3.3.2.2 GoC responses

Following punctuate sensory stimulation *in vivo*, GoCs have been reported to respond with short bursts of 2–3 spikes at up to 200–300 Hz (Vos et al., 1999). In BSB simulations, GoCs immersed in the GrC active cluster generated a burst of 2-5 spikes with a maximum instantaneous frequency of 213 ± 29 Hz (Fig. 4c). When GABA synapses and gap junctions between GoCs were switched off, the response bursts showed up to 6 spikes, with a higher maximum instantaneous frequency (308 ± 16 Hz) (n=70 GoCs; p < 0.001, paired *t*-test) (Fig. 4c). The burst was caused by synaptic excitation relayed by Gloms and GrCs (through both *aa*s and *pf*s), which generated AMPA and NMDA currents in GoC dendrites (Movie S2). The “silent pause” appearing after the burst was caused both by an intrinsic phase-reset mechanism (Holtzman et al., 2006; Solinas et al., 2006; Vos et al., 1999) and by reciprocal inhibition between GoCs, demonstrating marked dendritic processing capabilities (Masoli et al., 2020a).

##### 3.3.2.3 PC and MLI responses

PCs *in vivo* are known to respond to punctuate stimulation with burst-pause patterns (Herzfeld et al., 2015; Ramakrishnan et al., 2016). In BSB simulations, PC responses depended on cell position relative to the *mf* active bundle (Fig. 5a). The PCs placed vertically on top of the GrC active cluster received the largest number of *aa* and *pf* synaptic inputs producing typical burst-pause patterns (Masoli and D’Angelo, 2017). The *burst coefficient* was correlated with the number of synaptic inputs from *pf* and *aa* (multivariate regression analysis: R^2^=0.91) (Fig. 5b). The *pause coefficient* (was correlated with the burst coefficient (NMI=0.79) and with the number of spikes from MLIs (NMI=0.66) (Fig. 5a), reflecting the origin of the pause from both intrinsic after-hyperpolarizing mechanisms and MLI inhibition (Masoli et al., 2017). Indeed, MLIs are known to narrow the time window and reduce the intensity of PC responses (Kim and Augustine, 2020). In BSB simulations, the PC AMPA current arose soon after the spikes emitted by GrCs, while the PC GABA current was delayed by 2.6 ms (Fig. 5c). In summary, the di-synaptic IPSCs produced by MLIs quickly counteracted the monosynaptic EPSCs produced by *aa*s and *pf*s, providing precise time control over PC activation (Bower and Woolston, 1983; Diwakar et al., 2011).

**Figure 5.**
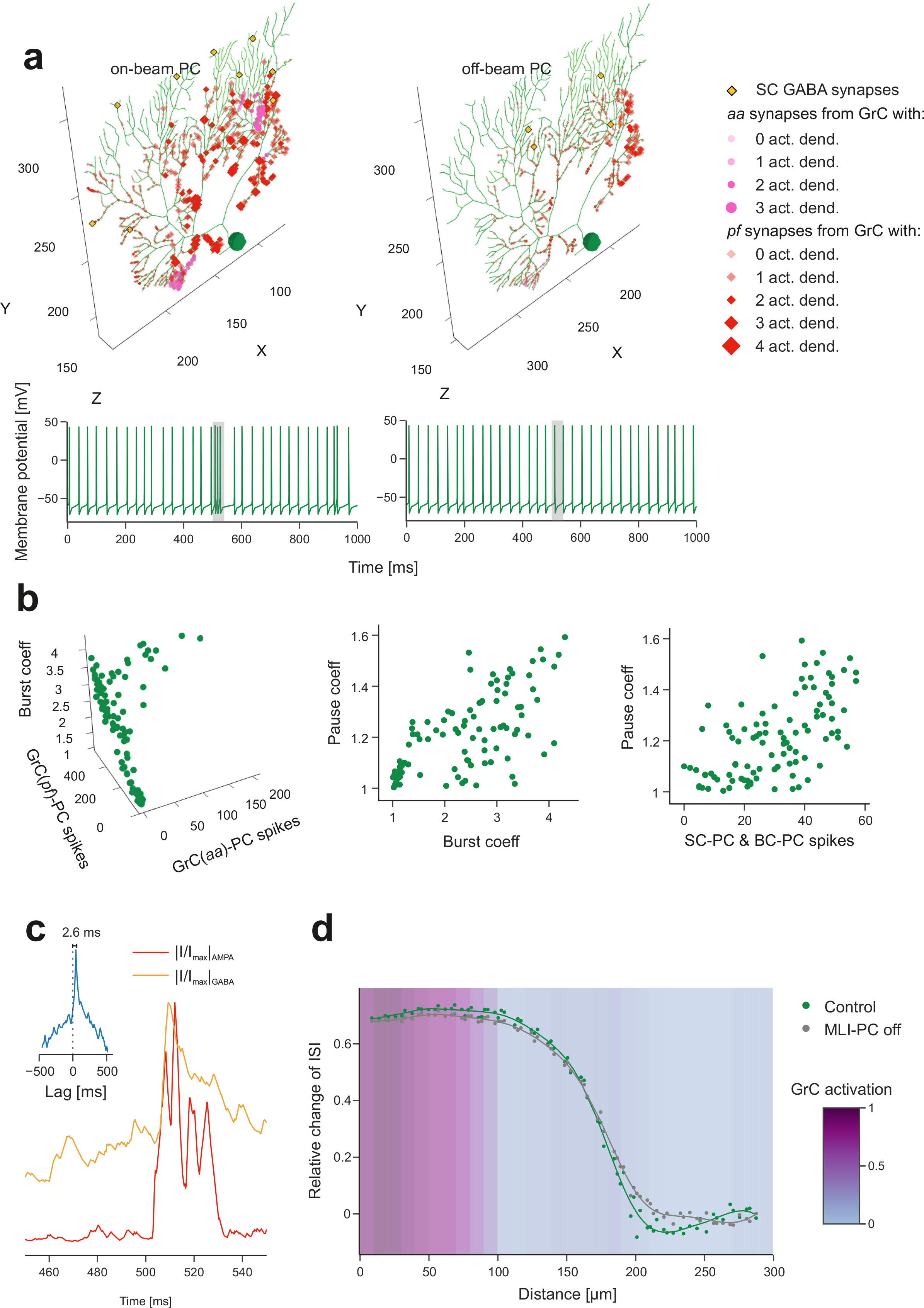
Purkinje cell activation. **(a)** Analysis of the burst-pause response of PCs to burst stimulation on 4 *mf*s (20ms@200Hz) over 4Hz background noise on all *mf*s. The *burst coefficient* (i.e. the shortening of the inter-spike interval due to the *mf* burst, with respect to baseline) is reported against the number of spikes from *aa*s and from *pf*s (multivariate regression analysis: R^2^=0.91). The *pause coefficient* (i.e. the elongation of the inter-spike interval after the *mf* burst response, with respect to baseline) is reported against either the *burst coefficient (*NMI=0.79) or the number of spikes from SCs and BCs (NMI=0.66). **(b)** The PC placed on top of the GrC active cluster and the PC placed at its margin show different synaptic inputs. GABAergic synapses from SCs are on medium-thickness dendrites (those from BCs on PC soma are not shown), *aa* synapses are located on thin dendrites and *pf* synapses on thick dendrites. In this example, the on-beam PC receives 23% of its *aa* synapses and 6% of its *pf* synapses from GrCs with at least 2 activated dendrites, the off-beam PC 0% of its *aa* synapses and 0.6% of its *pf* synapses from GrCs with at least 2 activated dendrites. The corresponding membrane potential traces are shown at the bottom (*mf* burst is highlighted by grey band). **(c)** Synaptic currents recorded from the PC on top of the GrC active cluster (same as in (b)), in voltage-clamp. The traces are the sum of all excitatory (AMPA) and inhibitory (GABA) dendritic currents during the *mf* burst. They are rectified, normalized and cross-correlated (inset) unveiling a GABA current lag of 2.6 ms with respect to AMPA current. **(d)** By stimulating a *mf* bundle (100ms@50 Hz Poisson stimulation on 24 adjacent *mf*s), the PC response modulation was quantified by the relative change of Inter-Spike-Interval (ISI), during the stimulus, where 0 corresponds to baseline. The two series of points compare PC response modulation when SCs and BCs were either connected or disconnected from PCs. The curves are regression fittings to the points (Kernel Ridge Regression using a radial basis function pairwise kernel, from Python scikit-learn library). The GrC active cluster was identified by a threshold on the stimulation-induced activity by using kernel density estimation.

BCs *in vivo* are known to generate lateral inhibition reducing PC discharge below baseline causing contrast enhancement (Eccles et al., 1967; Kim and Augustine, 2020). In BSB simulations, this pattern emerged during stimulation of a *mf* bundle (100 ms @ 50 Hz stimulation on 24 neighboring *mf*s). The PCs placed in a stripe 150-200 μm beside the active cluster along the transverse axis were inhibited bringing their frequency below baseline. When MLIs-PC synapses were switched off, the effect disappeared (Fig. 5d) revealing contrast enhancement due to lateral inhibition.

The response of MLIs *in vivo* is only partially known (Eccles et al., 1967). In BSB simulations, SCs and BCs intersected by active *pf*s responded to input bursts and their activity remained higher than baseline for several hundreds of milliseconds, especially in SCs (Rizza et al., 2021) (Fig. S4b).

## 4 Discussion

This work develops and benchmarks a new framework for multi-scale brain modelling (BSB), which accounts for the richness of network architectures and the variety of dynamic processes generating neural activity at different spatial and temporal scales. Although effective tools for microcircuit modelling appeared recently [BMTK (Billeh et al., 2020; Dai et al., 2020), NetPyNE (Dura-Bernal et al., 2019), Snudda (Hjorth et al., 2020) PyCabnn (Wichert et al., 2020)], their connectivity rules deal well with population-level and probabilistic approaches but a subset of modelling problems remains unsolved, when it comes to dealing with neurons as entities in space with specific morphologies. BSB addresses these needs with a set of tools designed to work with complex network topologies, cell morphologies and many other spatial and *n*-point problems. These properties allow BSB to fully empower a “scaffold” modeling strategy, in which a specific brain region or cell type can be modelled and specific cell placement or connectivity datasets can be changed without having to regenerate the entire network. An extended comparison of BSB with other network modeling tools is shown in Fig. S1.

BSB leveraged on a handful of fundamental data (cell density and placement) and on neuronal morphologies hosting accurate biophysical models and, by means of geometry-based connectivity rules, generated a new ground-truth for the cerebellar cortical network. BSB extracted information from the interdependence of parameters generating the connectome and allowed us to fill gaps in knowledge through constructive rules providing testable predictions on network dynamics. For the first time, functional simulations using naturalistic stimuli revealed the spatio-temporal dynamics across the entire cerebellar microcircuit at cellular resolution (Fig. 6) (Casali et al., 2020).

**Figure 6.**
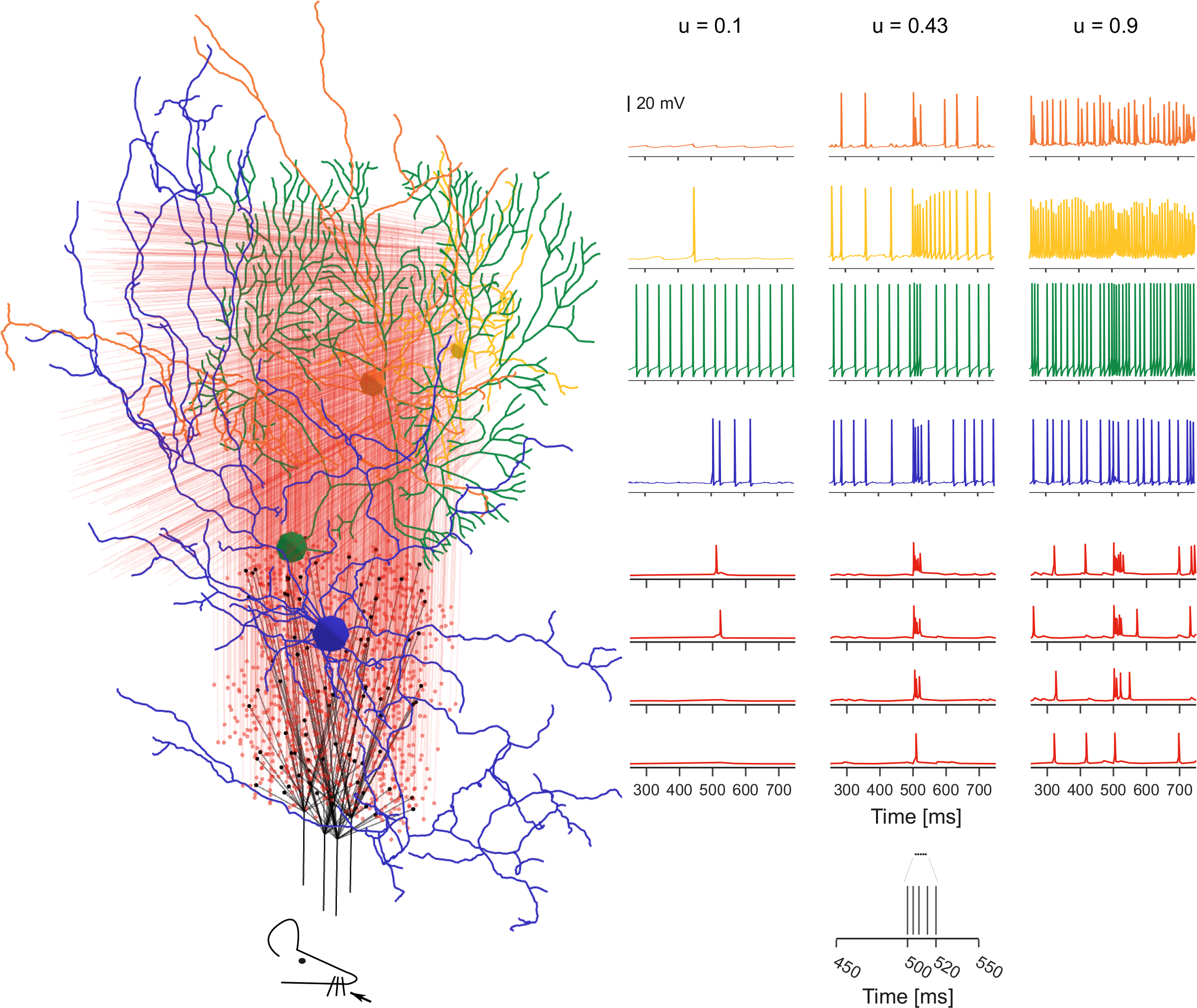
Activation of a vertical column of the cerebellar cortex. The whisker air-puff stimulus is delivered to 4 adjacent *mf*s, which branch in 4 Glom clusters. GrCs responded rapidly with a burst when at least 2 dendrites were activated. A GrC dense cluster is formed and the signal propagates up through an *aa* bundle and transversally along a *pf* beam. GoCs receive the signal both on basal and apical dendrites. PCs vertically on top of the active cluster are invested by *aa* and *pf* synaptic inputs. On-beam SCs and BCs receive signal through *pf* synapses; SC axons inhibit mainly on-beam PCs, while BC axons inhibit mainly off-beam PCs. The membrane potential traces (*mf* burst starts at 500 ms) are shown for each neuronal population. Traces in the three columns correspond to three different release probabilities at the *mf*-GrC synapses: u=0.1, u=0.43 (control condition used in the rest of the paper), u=0.9. The lower and higher u-values are typical of long-term synaptic depression and potentiation reported in the granular layer.

Geometrical rules deriving from anatomical and physiological works (Billings et al., 2014; D’Angelo et al., 2016) almost completely anticipated network connectivity at the cerebellar input stage. Each Glom hosted ∼50 excitatory and ∼50 inhibitory synapses on as many GrC dendrites, plus ∼2 excitatory synapse on basolateral dendrites from as many 2 GoCs, summing up to ∼102 synapses per Glom, in agreement with the anatomical estimate of ∼200 (Hamori et al., 1997). Each one of the 4 GrC dendrites received an excitatory and (in most cases) an inhibitory input from as many different *mf*s and GoCs (Mapelli et al., 2014; Tabuchi et al., 2019). Each GoC received ∼320 *aa* synapses on basal dendrites and ∼910 *pf* synapses on apical dendrites (Cesana et al., 2013) and there were ∼2 electrical synapses per GoC-GoC pair (Dugué et al., 2009). Only the number of GoC-GoC GABAergic synapses, which amounted to a figure of 160 after functional tuning, lacked an experimental counterpart.

In the molecular layer, under geometric and functional constraints, the BSB model placed limits to the debated numbers determining PC and MLI connectivity. The model predicted that ∼25% of *aas* contacted the distal dendrites of the overlaying PCs (7’133 out of 28’615 GrCs), each *aa* forming 2.4 synapses on average, supporting the important role of *aa*s (Bower and Woolston, 1983; Lu et al., 2009). And since *pfs* formed 1 synapse per PC dendritic intersection, each PC received 12% of the whole GrC inputs from *aa*s, matching the empirical estimate of 7-24% (Gundappa-Sulur et al., 1999). The BSB model generated ∼25 SC-PC and BC-PC synapses altogether, which compares well with the experimental estimate of ∼20 (Lennon et al., 2014). Moreover, there were ∼17’600 *pf*-MLI-PC synapses (∼2’600 *pf*-SC-PC and ∼15’000 *pf*-BC-PC synapses), compatible with the prediction that the *pf* -MLIs-PC input is larger than the *pf-PC* input on the same PC (Jörntell et al., 2010). In general, since all dendritic trees in the molecular layer are orthogonal to *pf*s, the BSB reconstruction ranked the number of synapses according to dendritic size - PC (∼1’500) > GoC (∼900) > BC (∼700) > SC (∼500) – a figure that would increase proportionately by scaling the model slab to include full-length *pfs* (∼3 mm) (Soha et al., 1997).

In aggregate, the BSB model reconstruction shows that geometrical organization largely determines cerebellar network connectivity, supporting the original intuition (Eccles et al., 1967; Marr, 1969). A similar conclusion was recently reported for the cortical microcolumn (Markram et al., 2015). Moreover, accurate single neuron models proved critical to carry out simulations allowing us to finetune the connectome. In particular, the number of inhibitory synapses per GoC-GoC pair was increased in order to make them fire at ∼19 Hz [2-30 Hz range: (Vos et al., 1999; Wilms and Häusser, 2015)]. Similarly, the number of inhibitory synapses per SC-SC and per BC-BC pairs was tuned in order to make them fire at ∼10 Hz [1–35 Hz range: (Jirenhed et al., 2013; Kim and Augustine, 2020)] and to bring PCs into their resting-state frequency range of ∼31 Hz [36.4 ± 11.5 Hz in : (Arancillo et al., 2015; Zhou et al., 2015) *in vivo*. While the connectivity between pairs of inhibitory interneurons warrants future experimental investigation, this BSB connectome ensured that GrCs and GoCs were entrained into low-frequency coherent oscillations in resting-state (Hartmann and Bower, 1998) under gap junction control (Dugué et al., 2009), providing high-level validation to the whole construct.

The functional validation of cerebellar network model was completed by simulating responses to naturalistic *mf* bursts, which rapidly propagated through the Grc-PC neuronal chain supporting the hypotheses of a cerebellar vertical organization (Bower and Woolston, 1983) and of burst-burst (or detonator) transmission (Rancz et al., 2007) (Fig. 6) (Movie S1). GrCs responded in dense clusters (Diwakar et al., 2011) regulated by GoCs, while PCs generated spots of activity regulated by MLIs (Movie S1). Not unexpectedly, SCs and BCs effectively reduced activation of PCs placed either along or beside the active *pf*s, respectively, generating feedforward and lateral inhibition. PCs showed the typical burst-pause responses that are thought to correlate with cerebellar-dependent behaviors (Herzfeld et al., 2015). These response patterns were seriously altered by changing microscopic parameters, such as *mf* neurotransmitter release probability (Masoli et al., 2020), whose effect propagated from the cerebellar input stage through the whole cortical network.

In conclusion, BSB allowed us to bind structure to function and dynamics and to demonstrate, for the first time, activity patterns in a vertical column of the cerebellar cortex. In the model, molecular and cellular properties reverberated across scales controlling spike timing and distribution. Given the BSB “scaffold” design, new neurons and cellular mechanisms could be easily plugged-in to update the model and apply it to investigations along ontogenesis, across species (for example in humans) and in pathology. The model may be used in the future to explain how genetic or epigenetic modifications would change PC responses at the cerebellar output, as it is supposed to happen in ataxia, dystonia and autism (Huang et al., 2021; Peter et al., 2016). The model may also be used to predict the impact of drugs acting on ionic channels and synaptic receptors. Finally, the models could host simplified neurons and run on hardware accelerators to implement large-scale simulators (e.g. Fig S6) and drive closed-loop controllers with learning inside neuromorphic computers and neuro-robots. In principle, the BSB workflow may be applied to any neuronal circuit promoting an ecosystem of modelling packages compatible with one another, for long-term value and extended use of brain modelling in the scientific community.

## 5 Methods

BSB is a Python (RRID:SCR_008394, version 3.6+) package that can be installed from *pip.* It is a proper Python library with a command line interface for the most common operations. It includes workflows and building tools for multiscale modelling of networks (both reconstruction and simulation) and is compatible with a wide variety of target systems such as personal computers, clusters or supercomputers and provides effortless parallelization using MPI.

### 5.1 The BSB modelling framework

There are 3 main phases in the scaffold modelling workflow that can be visited iteratively when changes need to be made: configuration, reconstruction and simulation. The core concepts of the framework during the reconstruction phase are i) the network *volume*, with the definition of various partitions such as layers, meshes or voxel sets (from brain atlases) and arranging elements which can be structured hierarchically to give rise a complex description of the entire region under consideration, ii) the *cell types* which determine the properties of cell populations, such as their spatial representation (soma radius, geometrical extension and/or morphologies), iii) the *placement* of said cell types into subspaces of the network volume using certain *placement strategies,* and iv) the c*onnectivity* between cell types using certain *connection strategies* With this information the framework places and connects the cells, storing the result in a network reconstruction file. Then the simulation phase follows, where *simulations* can be defined with *cell models, synapse models* and *devices* which together define the simulator specific representation of respectively cell types, connection types and in/output variables.

All the above concepts (Fig. 1) can be defined in a preceding configuration phase, either in a configuration file and declared programmatically. The framework was developed with a modular architecture in mind, where each module revolves around a central polymorphic class: the placement module has its *PlacementStrategy* and the connectivity module its *ConnectionStrategy*. The users can provide implementations of any interface to extend the repertoire of default placement or connectivity strategies with their own. Each user-defined strategy has access to its configuration node, so users can flexibly configure and parametrize their strategies, leveraging the support provided by the framework.

Various other interfaces exist for less commonly extended functions of the framework, such as configuration parsers (a JSON parser is provided), simulator backends, storage engines (HDF5 and SONATA backends are provided) or CLI commands.

The scaffold builder compiles its models into an HDF5 file or SONATA files, a format standard for neural models proposed by the Blue Brain Project and Allen Institute for Brain Science (Dai et al., 2019). The HDF5 file is a self-contained hierarchical file that includes all the required information to deploy the model on another machine, including configuration, placement, connectivity and even morphology information. These network architecture files can then be used to reproduce results, optimize parameters or to run entirely different simulations using the same structural information. Parts of the reconstruction can be repeated independently, and the datasets overwritten or appended with the results. This allows for quick tentative changes to be made, which improves iteration times of model development, specific cases and parameter exploration.

#### 5.1.1 Placement

The placement is organized into placement objects that consider certain cell types, and a subspace of the volume. These objects determine the number, position, morphology and orientation of each cell, according to the desired placement strategy. A variety of configuration mechanisms exist to define the number of elements to be placed, such as a fixed count, a specific density (volumetric or planar) or a ratio to the density of another type. Other elements can be instantiated as well, with or without 3D positions for other purposes (e.g., fibers with their somatic origin outside the considered brain circuit). A post-processing step after placement may be enabled, where the elements can be pruned, moved, or labelled (e.g., labelling separate zones with their own connectivity patterns or identifying individuals to be hubs in a modular network). Each morphology can be rotated based on the voxel orientation in which it is placed, and fibers crossing multiple voxels can be bended, in order to follow the detailed folding of the region.

The main provided placement strategies are (Fig. 1b):

##### Particle placement

The neurons are placed randomly and then checked for collisions, using kd-tree partitioning of the 3D space (Bentley, 1975). Colliding particles repel each other until they no longer collide, where the inertia of the particle is proportional to the radius. It is computationally efficient, yields uniform placement in 3D space, working properly even in irregular shapes, and it can deal with multiple cell types of different size. A pruning step can be enabled to remove cells positioned outside the desired subspace.

##### Parallel array placement

The neurons are placed in parallel rows on a desired surface, with a certain angle and specific distances between adjacent cells. A direction-specific jitter can be configured.

##### Satellite placement

The neurons are placed near each cell of an associated type (planet cells). Satellite positions are chosen at a random distance within a range based on the radii of the associated cell types, so that each planet cell has a certain number of satellite cells around it (Casali et al., 2019).

#### 5.1.2 Connectivity

Each connection identifies a pre-synaptic element and a post-synaptic element. When multi-compartmental neuron models are used, the synaptic locations on specific morphology compartments are also identified. Connections may target either populations, subpopulations, or only specific regions of the cell morphologies. A post-processing step after connectivity may be enabled, where the identified synapse locations can be re-distributed (e.g. pruning or moving the synapses). The use of cell morphologies can be combined with soma-only approaches. Multiple synapses per pair can be requested, following a probability distribution.

The main provided connectivity strategies are (Fig. 1c):

##### Touch detection

The 3D space is partitioned using a kd-tree to search for potentially intersecting cell pairs. Then, the actual points of intersection are determined using another kd-tree specific to the cell pair morphologies, with a maximum distance parameter.

##### Voxel intersection

Each pre-synaptic cell is represented by a voxelized morphology and these voxels are tested for intersections with the voxels of the postsynaptic cells, using R-tree 3D space partitioning. When matching voxels are found, random compartments in each voxel are selected, introducing variability. It is, in this respect, less deterministic than the touch detection strategy (Nolte et al., 2019). A *FiberIntersection* variant exists to optimize the case of long, thin neurites, whose path can be deformed through space according to a 3D field of direction vectors.

##### Distance-based in/out degree

When only the distance distribution is given, cells connection probability is based on their distance. When network topology matters, the distance-based probability can be combined with indegree and outdegree distributions, the number of postsynaptic elements per presynaptic element is determined by samples of the outdegree distribution. During sampling each postsynaptic target is weighted according to their distance probability and their probability to transition from their current indegree N to indegree N+1, as dictated by the indegree distribution. To optimize the algorithm, a kd-tree is queried for cells within a maximum search radius derived from the cumulative distance probability.

#### 5.1.3 Simulation

BSB can instruct simulators to run the configured models. Although multiple adapters to different simulators are provided (**NEURON** - RRID:SCR_005393, **NEST** - RRID:SCR_002963, **Arbor** - 10.5281/zenodo.4428108), there is no common high-level language to send instructions across simulators. Instead, sets of simulator-specific configuration expose the simulators’ underlying APIs more directly. These classes contain the simulator specific logic to fully prepare a simulator to define inputs, execute, monitor progress, and collect output of simulations. The interface to the NEURON simulator has been applied specifically in this work.

##### NEURON adapter

We provide a simulator adapter for NEURON (Hines and Carnevale, 2001) that cooperates with our Python packages: *Arborize* to create high-level descriptions of cell models [https://github.com/dbbs-lab/arborize], *Patch* to provide a convenience layer on top of NEURON [https://github.com/Helveg/patch], and *Glia* to manage NMODL file dependencies and versioning [https://github.com/dbbs-lab/glia]. Together, these packages and the NEURON adapter provide out-of-the-box load balanced parallel simulations in NEURON. The adapter is capable of creating and connecting these *Arborized* cell models over multiple cores, implements a few basic device models such as spike generators, voltage and synapse recorders and collects the requested measurements in an HDF5 results file. The recorders can specify targets at the cellular or subcellular level, recording membrane or synapse voltages, conductances, currents and ionic concentrations. These easy configurable devices allow to monitor all signals propagating across the network to reproduce results at multiple scales, all tasks that usually require a considerable amount of effort in NEURON simulations.

#### 5.1.4 Visualization

BSB provides a plotting module to directly visualize simulation results including 3D network plots, cell activity in 3D space, PSTH, raster plots and more. BSB provides a Blender module containing a complete blender pipeline for rendering videos of the network activity on a single machine or a cluster. It can animate simulation results or be used to troubleshoot the model when BSB is used inside of Blender Python scripts. Functions exist to synchronize the state of the network with the Blender scene, to animate results or to generate *debug frames* to animate and troubleshoot placement, connectivity or simulation issues.

### 5.2 Cerebellar use case

Using this framework, a mouse cerebellar cortex microcircuit was reconstructed and simulated to benchmark and test the entire workflow. The example here reported refers to a volume partitioned in granular, Purkinje and molecular layer. Specifically, volume extended 300 μm along x, 200 μm along z, 295 μm along y (layers thickness: 130 μm granular layer, 15 μm Purkinje cell layer, 150 μm molecular layer). In the reference system, x-y is the sagittal plane, x-z the axial plane, z-y the coronal plane. The reconstructed volume was overall 17.7·10^-3^ mm^3^. The model was filled with biophysically detailed compartmental neurons for each cell type. Some structural data and multiple observations from electrical recordings *in vivo* and *in vitro* were used as constraints in building the model, further experimental measurements were used for structural and functional validation. Moreover, a version using extended generalized leaky integrate and fire point-neuron models (Geminiani et al., 2018) was generated and tested by NEST adapter (shown in Supplementary Material – Fig. S4).

#### 5.2.1 Cerebellar neuron placement

Both the granular layer and the molecular layer were filled by using *particle placement*. The granular layer is made up of densely packed granule cells (GrC) and glomeruli (Glom) intercalated with Golgi cells (GoC). Furthermore, a certain number of mossy fibers (*mf*) was created (without any 3D position), each terminating in about 20 glomeruli. Each GrC emits an ascending axon (*aa*) that raises perpendicularly to the overlying cerebellar surface and reaches the molecular layer bifurcating into two opposite branches of a parallel fiber (*pf*) elongating on the z-axis (major lamellar axis). The molecular layer was divided into a superficial sublayer (2/3 of the thickness) hosting the stellate cells (SC) and a deep sublayer (1/3 of the thickness) hosting the basket cells (BC) (Sultan and Bower, 1998; Whitney et al., 2009).

The Purkinje cells (PC) were placed on an axial plane (x-z) by using *parallel array placement.* PCs were placed along parallel lines, with an inclination angle of about 70° with respect to the z-axis (major lamellar axis). The dendritic tree of the PC is flattened on the sagittal plane and extends for about 150 µm (Masoli and D’Angelo, 2017). The parallel lines were defined at such a distance that the PC dendritic trees did not overlap, while along the major lamellar axis their somata could be packed closely together. For each neuronal population, the nearest neighbour, the pairwise distance distribution, and the radial distribution function were computed.

#### 5.2.2 Cerebellar microcircuit connectivity

The connectome of the cerebellar network took into consideration 16 connection types (identified by their source and target neuronal population): *mf*-Glom, Glom-GrC, Glom-GoC, GoC-GrC, GoC-GoC, GrC (*aa*)-GoC, GrC (*aa*) -PC, GrC (*pf*)-GoC, GrC (*pf*)-PC, GrC (*pf*)-SC, GrC (*pf*)-BC,SC-PC, BC-PC, SC-SC, BC-BC, GoC-GoC, GoC-GoC gap junctions.

Glomerular connectivity is a special case since it is largely constrained by prescribed neuroanatomical and neurophysiological information. Then, since the Gloms did not have defined morphological model, they were connected through probability strategies (*distance-based in/out degree*) to identify nearby compartments for synaptic locations on the target cell types, for which a realistic morphology was used. Specifically:

##### *mf*-Glom

The *mf* arborization creates anisotropic clusters of glomeruli and clusters originating from different *mfs* mixed up with each other to some degree (Sultan, 2001). Taking into account short branches since the small reconstructed volume, a local branching algorithm grouped glomeruli (a normal distribution of 20 ± 3) receiving signals from the same *mf* by a distance-based probability rule.

##### Glom-GrC

For each GrC, a pool of nearby Gloms were selected based on the distance between the Glom barycenter and the GrC soma center, and a maximum extension of GrC dendrites of 40 µm (Houston et al., 2017). From the pool, 4 Gloms, each from a different cluster, were randomly sampled and connected with one of the 4 GrC dendrites.

##### Glom-GoC

For each Glom, all GoCs with their soma at a radial distance less than 50 µm (corresponding to an average extension of GoC basolateral dendrites, isotropically in 3D (Cesana et al., 2013; Solinas et al., 2010)) were connected. The synapse was placed on a basal dendrite using an exponential distribution favouring the compartments closer to the centre of the glomerulus.

##### GoC-GrC

This connection absorbed the connection **GoC-Glom** which was generated using 3D proximity and a mean divergence (out-degree) of 40; then, each GoC synapsed directly on all GrCs that shared those glomeruli.

The rest of the cerebellar connectome was reconstructed by applying *voxel* and *fiber intersection,* and *touch detection*. *Voxel intersection* was preferred when 3D morphologies were intersecting. This strategy, by introducing a cubic convolution and randomization, reduced overfitting artifacts arising from the intersection of identical morphologies arranged in the quasi-crystalline cerebellar microcircuit. *Touch detection* and *fiber intersection* were preferred when dealing with 2D fibers (i.e. *aa* and *pf*), for which the *voxel intersection* would create cubic volumes not representative of the fiber as line segment. Therefore, for the connections involving *aa*s (GrC (*aa*)-GoC and GrC (*aa*) -PC), *touch detection* was applied with a tolerance distance of 3 µm. For the connections involving *pf*s (GrC (*pf*)-GoC, GrC (*pf*)-PC, GrC (*pf*)-SC, GrC (*pf*)-BC), *fiber intersection* was applied with affinity equal to 0.1 and a resolution of 20 µm. For the other connections, *voxel intersection* was applied: GoC-GoC, SC-SC, and BC-BC with affinity 0.5; SC-PC with affinity 0.1; BC-PC with affinity 1, GoC-GoC gap junctions with affinity 0.2. For chemical synapses (connection typologies from 1 to 15), the presynaptic compartment was always axonal and the postsynaptic compartment was dendritic or somatic (as for the BC-PC connection). For electrical synapses, gap junctions were created between dendrites. Following biological indications, specific sectors of morphologies were selected as source or target for synaptic localization, and, for each connected cell pair, the desired number of synapses was defined, eventually as a normal distribution (mean ± sd):

1. *mf*-Glom. Each *mf* spreads the input signal into a cluster of Gloms; no synapses are involved.
2. Glom-GrC. 1 excitatory synapse on the terminal compartment of each GrC dendrite.
3. Glom-GoC. 1 excitatory synapse on GoC basal dendrites.
4. GoC-GrC. 1 inhibitory synapse per GrC dendrite, on the pre-terminal compartment (Mapelli et al., 2009).
5. GoC-GoC: 160 ± 5 inhibitory synapses on GoC basal dendrites
6. GrC(*aa*)-GoC: 1 excitatory synapse on GoC basal dendrites
7. GrC(*aa*)-PC: 4 ± 0.4 excitatory synapses on PC dendrites with diameter < 0.75 μm (Masoli and D’Angelo, 2017; Wilms and Häusser, 2015)
8. GrC (*pf*)-GoC: 1 excitatory synapse on GoC apical dendrites
9. GrC (*pf*)-PC: 1 excitatory synapse on PC dendrites with diameter between 0.75 and 1.6 μm
10. GrC (*pf)*-SC: 1 excitatory synapse on SC dendrites
11. GrC (*pf*)-BC: 1 excitatory synapse on BC dendrites
12. SC-PC: 5 ± 0.5 inhibitory synapses on PC dendrites with diameter between 0.3 and 1.6 μm (Ango et al., 2008; Lu et al., 2009; Masoli et al., 2017)
13. BC-PC: 1 inhibitory synapse on PC soma
14. SC-SC: 100 ± 4 inhibitory synapses on SC dendrites
15. BC-BC: 100 ± 4 inhibitory synapses on BC dendrites
16. GoC-GoC gap junctions: 3 ± 1 gap junctions on GoC basal dendrites (symmetrically bidirectional) (Szoboszlay et al., 2016)

It is worth noting the use of specific subareas of the dendritic trees of GoC and PC as specific targets for synapse formation. GoCs receive inhibitory and electrical synapses from other GoCs and *aa* synapses on basal dendrites within granular layer, while *pf* synapses on apical dendrites within the molecular layer (Cesana et al., 2013). Concerning PCs, *aa* synaptic input projects onto a different part of the PC dendritic tree than *pf*, and SCs also target PC dendrites with thickness in a specific range (Lu et al., 2009; Masoli et al., 2017).

#### 5.2.3 Multi-compartmental neuron and synaptic models

Detailed multi-compartmental models of GrC, GoC, PC, SC, BC are available, in which dendritic and axonal processes are endowed with voltage-dependent ionic channels and synaptic receptors. In each model, cell-specific aspects critical for function are reproduced, e.g., the role of the axon initial segment, spontaneous firing and synaptic burst/pause behaviour. The following receptor-channel models, all validated against *in vitro* recordings, and generic gap-junction models were inserted in the appropriate neuron compartments:

- GrC synapses (Masoli et al., 2020b). mf-GrC: AMPA and NMDA receptors; GoC-GrC: GABAalpha1/6 receptors.
- GoC synapses (Masoli et al., 2020a). *pf*-GoC: AMPA; *aa*-GoC: AMPA and NMDA; *mf* -GoC: AMPA and NMDA; GoC-GoC: GABAalpha1, gap junctions (Szoboszlay et al., 2016; Vervaeke et al., 2012).
- PC (Z+ type) synapses(Masoli et al., 2017). *pf*-PC *and aa*-PC: AMPA; SC-PC and BC-PC: GABAalpha1.
- SC and BC synapses (Rizza et al., 2021). *pf* -SC and *pf* -BC: AMPA and NMDA; SC-SC and BC-BC: GABAalpha1.

Chemical neurotransmission was modelled using the Tzodyks and Markram scheme (Markram and Tsodyks, 1996; Tsodyks and Markram, 1997) on neurotransmitter release and receptors kinetic schemes for postsynaptic receptor activation. Glutamatergic neurotransmission could activate only AMPA either both AMPA and NMDA receptors (Nieus et al., 2006). GABAergic neurotransmission activated GABA-A receptors (Nieus et al., 2014). A neurotransmitter impulse was followed by a slow diffusion wave generating both a transient and a sustained component of the postsynaptic response, as observed experimentally. Parameters describing probability of release, diffusion, ionic receptor mechanisms, vesicle cycling parameters, time constant of recovery, electrical conduction were derived from original papers reporting GrC synapses (Diwakar et al., 2009; Masoli et al., 2020b), GoC synapses (Masoli et al., 2020a), GoC gap junctions (Szoboszlay et al., 2016), PC synapses (Masoli and D’Angelo, 2017), SC synapses (Rizza et al., 2021).

#### 5.2.4 Network simulations: stimulation and analysis

All simulations used the NEURON adapter of the BSB and were run in parallel through MPI on the CSCS *Piz Daint* supercomputer, with a time resolution of 0.025 ms. Simulations started with a 5-s stabilization period followed by a 100 ms initialization period, in which random *mf* inputs desynchronized the network. In all simulations, spikes and voltage traces at soma of all neurons were recorded. Depending on specific analyses, in some simulations further microscopic variables were recorded, as explained below. A set of stimulation protocols reproducing specific spatiotemporal patterns of *mf* activity was used to functionally validate the cerebellar network model; in some cases, the protocols were repeated using an altered version of the network model in terms of connectome (“by-lesion” approach), to quantitatively check the relative roles of the connectivity types.

##### Diffused background stimulation

The cerebellum *in vivo* is constantly bombarded by a diffused background noise, which determines the resting state activity of neurons and is thought to entrain the network into coherent low-frequency oscillations (Hartmann and Bower, 1998; Solinas et al., 2006; Vos et al., 1999). Therefore, we first explored the response of the network model to a random Poisson noise at 4 Hz (Rancz et al., 2007) on all *mf*s for 4 seconds, proving a testbench to validate the structural and functional network balance.

###### Steady state analysis

We compared basal discharges in the network to those recorded at rest *in vivo*. The mean frequency of each population was computed.

###### Oscillatory state analysis

We investigated the emergence of low-frequency coherent oscillations in the GoC and GrC populations. The power spectrum of GoC and GrC firing activity was computed by Fast Fourier Transform (FFT), applied to time-binned spike-counts (2.5 ms bins). The zero-component was cut off and the FFT was smoothed using a Savitzki-Golay filter (6th order polynomial, window of 51 bins). The same analysis was performed when GoC-GoC gap junctions were blocked, in order to check the role of electrical coupling in oscillatory behaviour of granular layer.

##### *mf* burst stimulation

The cerebellum *in vivo* responds with localized burst-burst patterns to facial or whisker sensory stimulation (Chadderton et al., 2004; Rancz et al., 2007). These bursts are supposed to run on collimated *mf* bundles generating dense response clusters in the granular layer and thereby activating the neuronal network downstream (Diwakar et al., 2009, 2011; Ramakrishnan et al., 2016; Roggeri et al., 2008). To simulate this functional response, we delivered a *mf* stimulus burst, superimposed on background noise at 4 Hz, to 4 *mf*s in the center of the axial plane, activating about 80 Gloms. The *mf* burst lasted 20 ms and was made of 5 spikes at fixed time instants (on average 200 Hz, maximum 250 Hz), within range of *in vivo* patterns (Chadderton et al., 2004; Rancz et al., 2007; Wilms and Häusser, 2015). Ten simulations were run to account for random variability of the background input and the network responses. Multiple variables over time were recorded: spike times and membrane voltages of every cell, synaptic currents in the dendrites of some cells, the internal Calcium ion concentration [Ca^2+^]_in_ in the dendrites of all GrCs and of some GoCs. Further simulations were carried out using different values of neurotransmitter release probability at the mf-GrC synapse (from u=0.43 to u=0.1 and to u=0.9).

###### General analysis of response patterns

For each neuronal population, a raster plot and a PSTH (peri-stimulus time histogram) was computed. Each population was described using a Multiple Regression Analysis: the dependent variable was the average firing frequency during 40 ms after the stimulus onset, over 10 simulations, the independent variables were the average numbers of spikes received from each presynaptic population. The linear regression was reported as direction coefficients and R^2^ score.

###### Analysis of granular layer responses

For GrCs we related the number of dendrites activated by the *mf* burst with the number of output spikes and the first-spike latency. The same protocol was carried out while switching-off phasic and tonic inhibition from GoCs (GABA-A receptor blockade). This allowed us to investigate excitatory-inhibitory loops in the granular layer, by estimating the response patterns of GrCs, their latencies, and the fraction of granule cells activated compared with the control condition. Furthermore, for each GrC, the level of [Ca^2+^]_in_ in the dendrites averaged on 500 ms from the *mf* burst was extracted and these [Ca^2+^]_in_ values were related to the number of dendrites activated by the stimulus. The correlation analysis was based on Normalized Mutual Information (NMI) (Laarne et al., 2021).

###### Analysis of PC responses

The PC response was analyzed to evaluate the burst-pause behavior. For each PC, an automatic algorithm extracted any shortening of the inter-spike intervals during the stimulus window (*burst coeff*.) and any elongation after the stimulus (*pause coeff*.) compared to baseline. The *burst coeff.* was correlated with the number of excitatory synaptic inputs (from *pf*s and *aa*s) by multivariate regression analysis. The *pause coeff*. was correlated with the number of inhibitory synaptic inputs (from MLIs) received during the burst stimulation (20 ms *mf* burst + 20 ms of delayed effects), by Normalized Mutual Information (NMI). Furthermore, the relation between the burst and pause coefficients themselves was analysed, by NMI. Further simulations were run clamping an on-beam PC at -70 mV, recording all synaptic currents. Excitatory synaptic currents (AMPA from *aa*s and *pf*s) and inhibitory synaptic currents (GABA from SCs and BCs) were summed up, then the cross-correlation among these two rectified and normalized currents was calculated, to identify the time lag. *Visualization of subcellular variables*. In some cases, ad-hoc computationally expensive recordings of multiple microscopic variables were performed. In an example focused on a GoC, all synaptic currents (AMPA and NMDA from Glom*s* on basal dendrites, AMPA and NMDA from *aa*s on basal dendrites, AMPA from *pf*s on apical dendrites, GABA from other GoCs on basal dendrites), gap junctions currents (from other GoCs on basal dendrites), and [Ca^2+^]_in_ were recorded and animated (see *Visualization*).

##### Lateral Poisson stimulation

The lateral inhibition from MLIs to PCs comes from activated MLIs providing inhibition to off-beam PCs, mainly from BCs due to their axon orientation (Kim and Augustine, 2020). To simulate this functional response, we delivered a 50 Hz Poisson distributed stimulus, lasting 100 ms, superimposed on the background noise (at 4 Hz), on 24 *mf*s on one side of the volume, to monitor the modulation of MLI inhibitory effects on PCs at different distances from the active cluster. Two conditions were evaluated: i) control and ii) MLIs disconnected from PCs. Ten simulations for each condition were carried out.

###### Analysis of PC responses

For each PC, the average Inter-Spike-Interval (ISI) during 200 ms baseline and the average Inter-Spike-Interval during the 100-ms stimulus was computed. The relationship between the distance of a PC from the active cluster and its activity modulation (balance between GrCs excitation and MLIs inhibition) was investigated in control condition and in the “no MLI-PC” condition.

## 6 Code availability

The BSB source code is available at https://github.com/dbbs-lab/bsb, and can be installed through *pip* as a Python package available at https://pypi.org/project/bsb/. Documentation can be found at https://bsb.readthedocs.io/.

## 8 Acknowledgements/funding

This research has received funding from the European Union’s Horizon 2020 Framework Program for Research and Innovation under the Specific Grant Agreement No. 945539 (Human Brain Project SGA3) and Specific Grant Agreement No. 785907 (Human Brain Project SGA2) and from Centro Fermi project “Local Neuronal Microcircuits” to ED. Special acknowledgement to EBRAINS and FENIX for informatic support and infrastructure.

## 9 Author contribution

CC and RDS designed and developed the informatic framework and performed the simulations; RDS wrote most of the code; AG, SM, MR, AA contributed with essential model components; RDS, CC, ED analyzed the data, wrote the manuscript, and prepared the figures; CC and ED coordinated the work and the EBRAINS interaction. ED promoted the project, supported it financially, defined the physiological aspects and finalized the manuscript.

## 11 Figure legends

Figures or their specific panels can be visualized as interactive *.html* files from https://dbbs-lab.github.io/deschepper-etal-2021/, adding e.g. “figure” + number + panel + “.html” e.g.:“figure2a.html”, on this URL.

## 12 Supplementary material

Supplementary figures and videos can be visualized as interactive *.html* files at https://dbbs-lab.github.io/deschepper-etal-2021/, adding e.g. “video” + number + “.html” e.g.: “videoS2.html”, on this URL.

**Figure S1.**
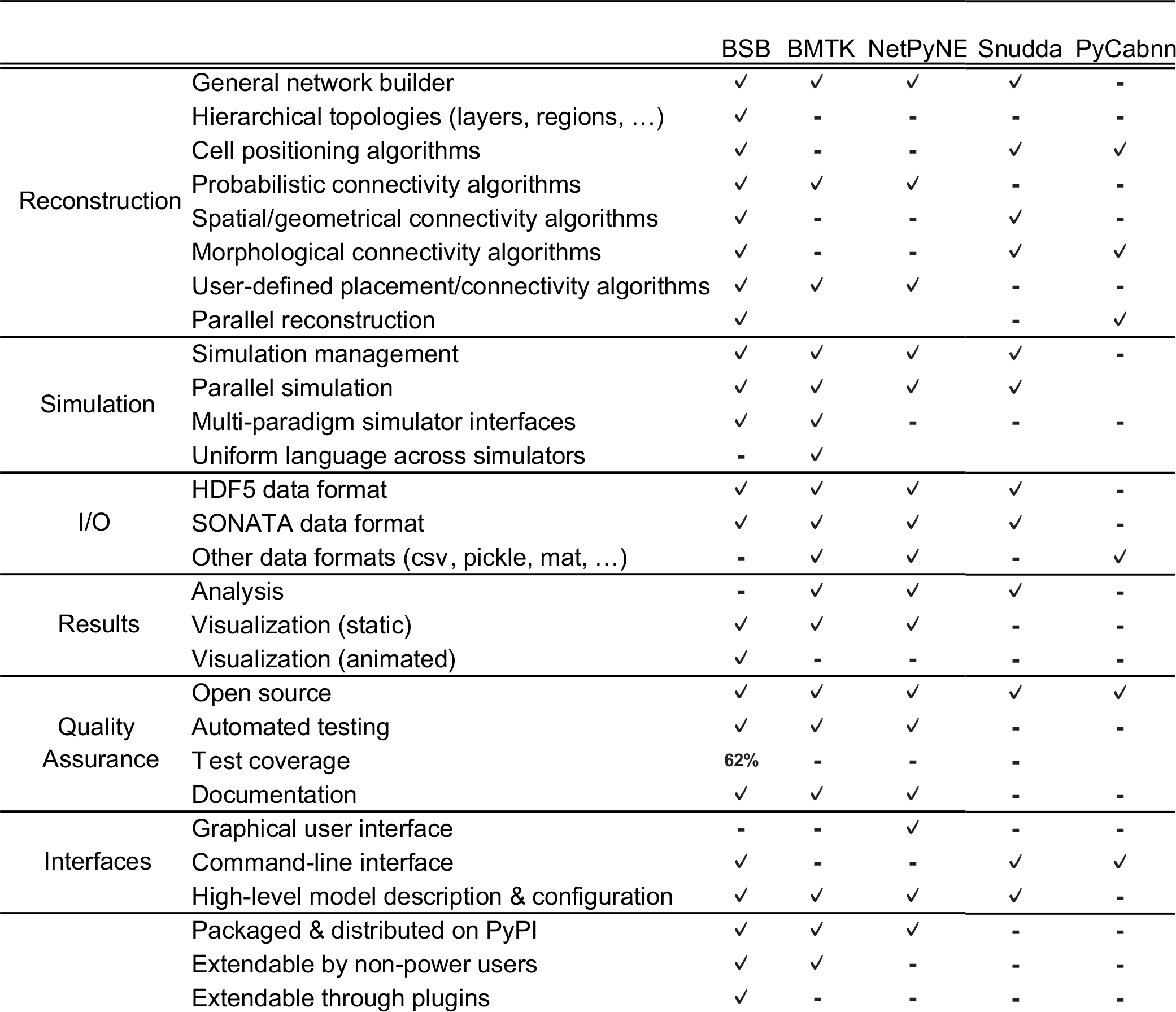
Modelling frameworks and tools. The table compares BSB at https://github.com/dbbs-lab/bsb https://pypi.org/project/bsb/ https://bsb.readthedocs.io/ with other brain modelling frameworks and specific-purpose tools, including Brain Modelling ToolKit (BMTK), NetPyNE, PyCabnn, Snudda:
https://github.com/AllenInstitute/bmtk.git https://github.com/Neurosim-lab/netpyne.git https://github.com/CNS-OIST/pycabnn https://github.com/Hjorthmedh/Snudda https://github.com/AllenInstitute/bmtk.git https://github.com/Neurosim-lab/netpyne.git https://github.com/CNS-OIST/pycabnn https://github.com/Hjorthmedh/Snudda The Allen Institute is developing the Brain Modelling ToolKit (BMTK) (Dai et al., 2020) that has been exploited to reconstruct cerebro-cortical microcircuits (Billeh et al., 2020) and provides an interface with multiple simulators. Another recent tool is NetPyNE (Dura-Bernal et al., 2019), with a graphical and Python interface for NEURON. NetPyne and BMTK also implement general modelling frameworks, with solutions to most common modelling problems and good useability. However, their connectivity rules deal well with population-level and probabilistic approaches, but a subset of modelling problems remains unsolved when it comes to dealing with neurons as entities in space with specific morphologies. BSB addressed these needs with a set of tools designed to work with complex network topologies, cell morphologies and many other spatial and *n*-point problems. Other tools are designed for specific purposes, e.g. Snudda supports random placement and touch detection only, and PyCabnn seems specifically targeted to the cerebellar granular layer. The main BSB advantages are (1) complex hierarchical descriptions of model topology (2) generalized strategies for cell placement and connectivity of either spherical somata or full neuron morphologies, (3) easily interchangeable components for placement/connectivity, cell models, synapse models (4) programming extensible to the core (plugins, interfaces, easy-to-override classes, everything is designed to be easily modified from outside by novice users with full integration in the configuration system), (5) organized workflow that removes boilerplate code and promotes stages that follow an intuitive modelling process, with a strict division between the model parameters in the configuration and code implementation, (6) integration with most of the available brain modelling toolkits, which have a Python interface and use standard data formats. These properties allow BSB to fully empower a “scaffold” modeling strategy, in which any brain region or cell type can be modelled, specific cell placement or connectivity datasets can be changed without having to regenerate the entire network, multiple configurations can be hosted. This strategy can drastically cut down on scientific resources and time required to investigate multiple network states. BSB functionalities can be explored by navigating the git-hub at https://bsb.readthedocs.io/en/latest/usage/getting-started.html.

**Figure S2.**
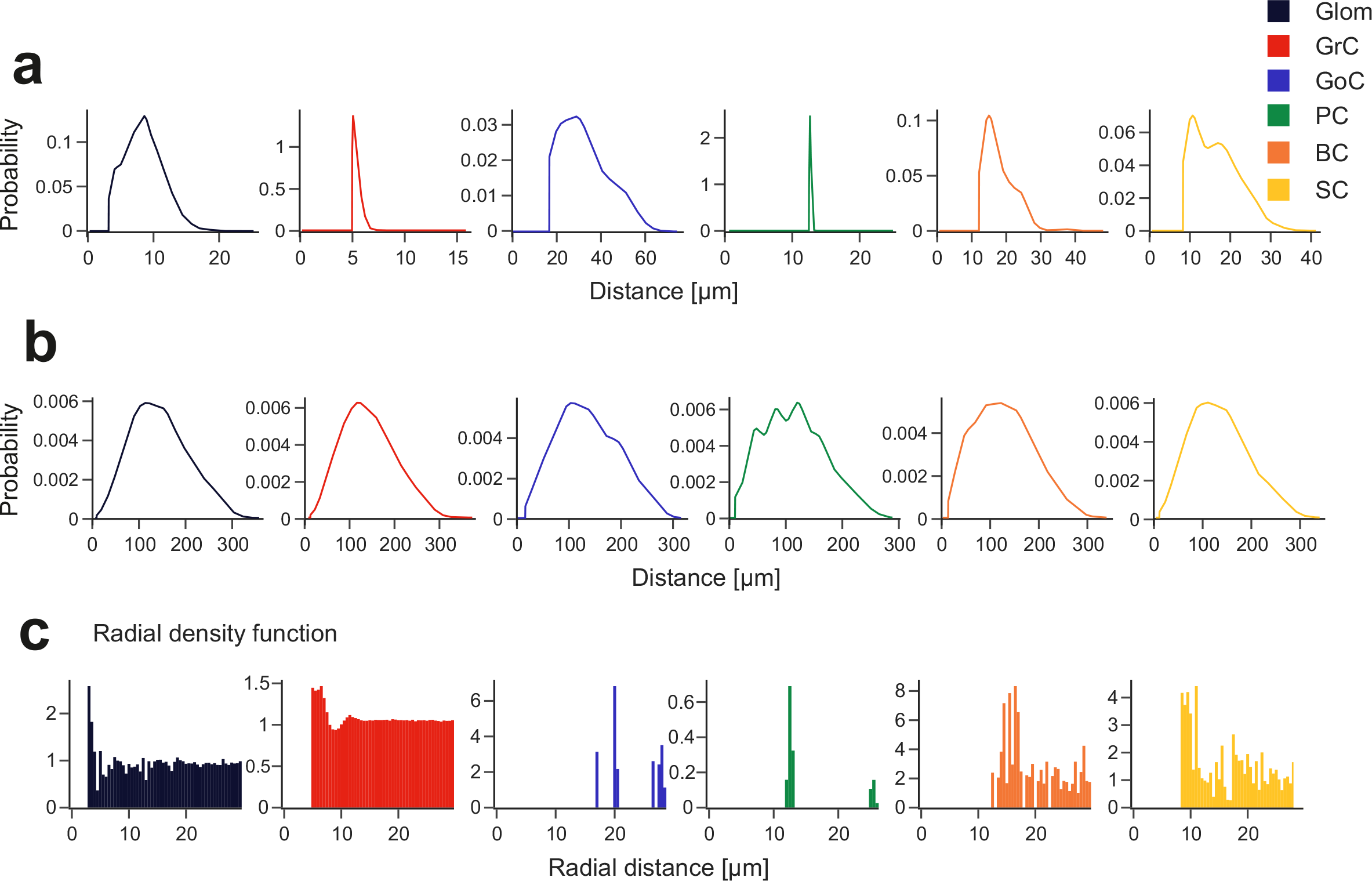
Placement metrics. Cell placement is assessed using various metrics for each neuron population including **(a)** Nearest Neighbor distance, **(b)** Pairwise Distance, **(c)** Radial Density Function. These metrics show realistic cell positioning without unphysiological lattice-like structures.

**Figure S3.**
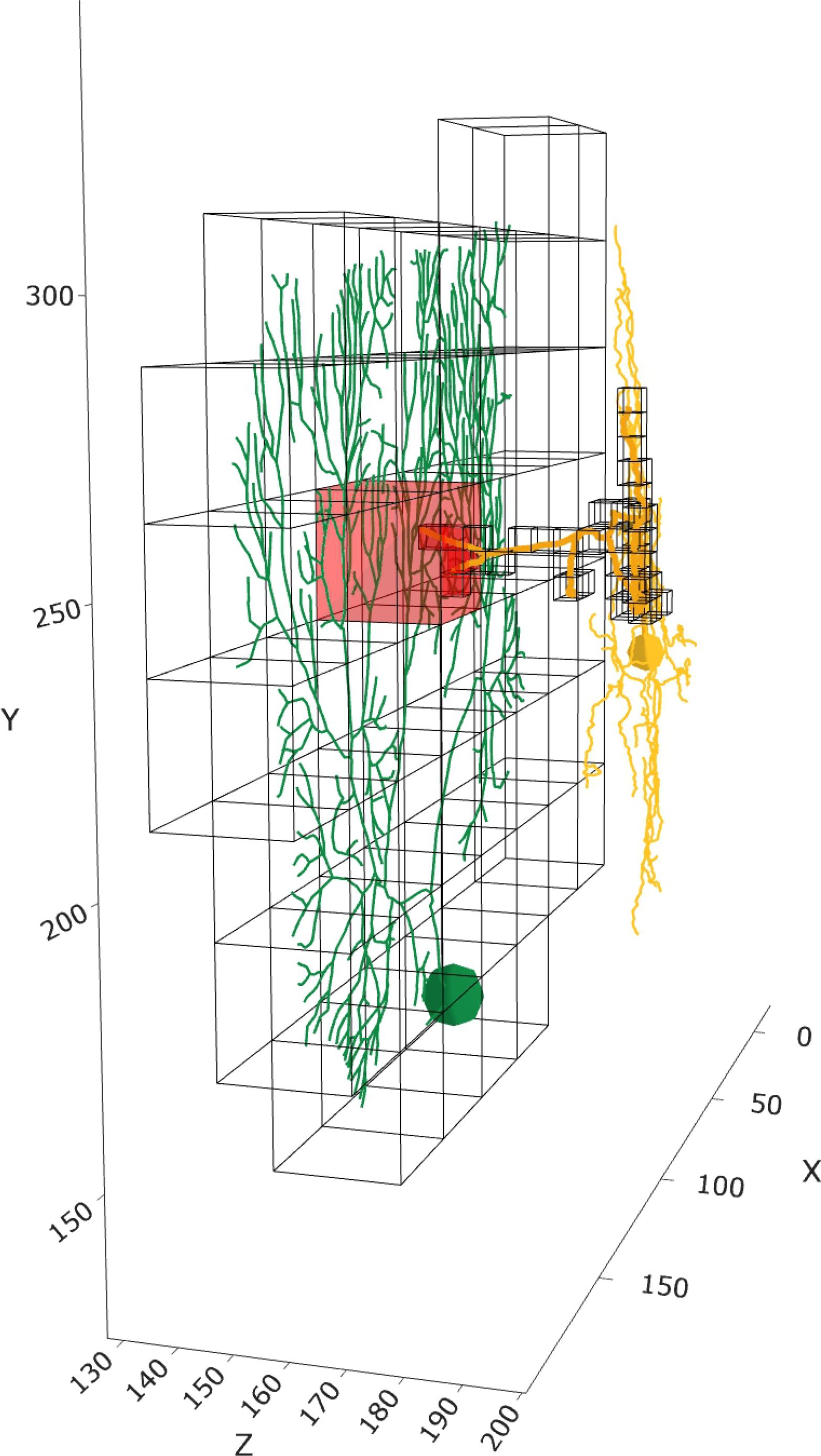
Connecting SC-PC by voxel intersection. A mesh of adjacent voxels is used to enwrap the axon of a stellate cell (50 cubes with 4.6 µm side) and the dendritic tree of a PC (50 cubes with 26 µm side). The intersecting voxels are in red. The synapses are located on compartments within the intersecting voxels.

**Figure S4.**
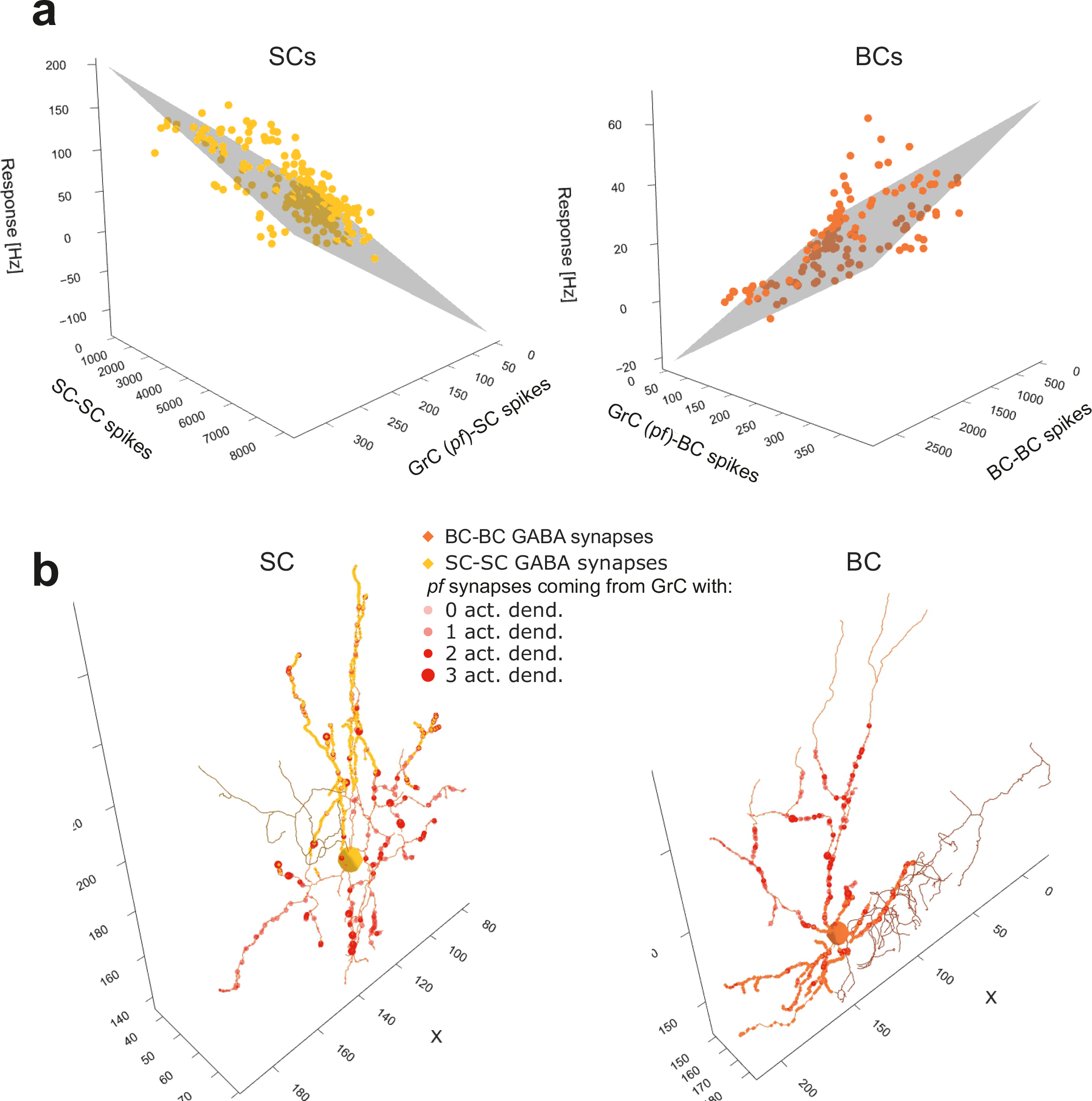
MLI responses to *mf* burst. **(a, b)** Multiple linear regression of SCs and BCs in responses to a *mf* burst (20ms@200Hz) against the number of synaptic spikes from *pf*s and from other SCs or BCs. **(c, d)** One SC and one BC crossed by an active *pf* beam are represented in 3D. The GABAergic synapses from other SCs or BCs are also indicated. Bigger markers correspond to presynaptic GrCs more activated by the *mf* burst. In this example, the SC receives 8% and the BC 7.5 % of their *pf* synapses from GrCs with at least 2 activated dendrites.

**Figure S5.**
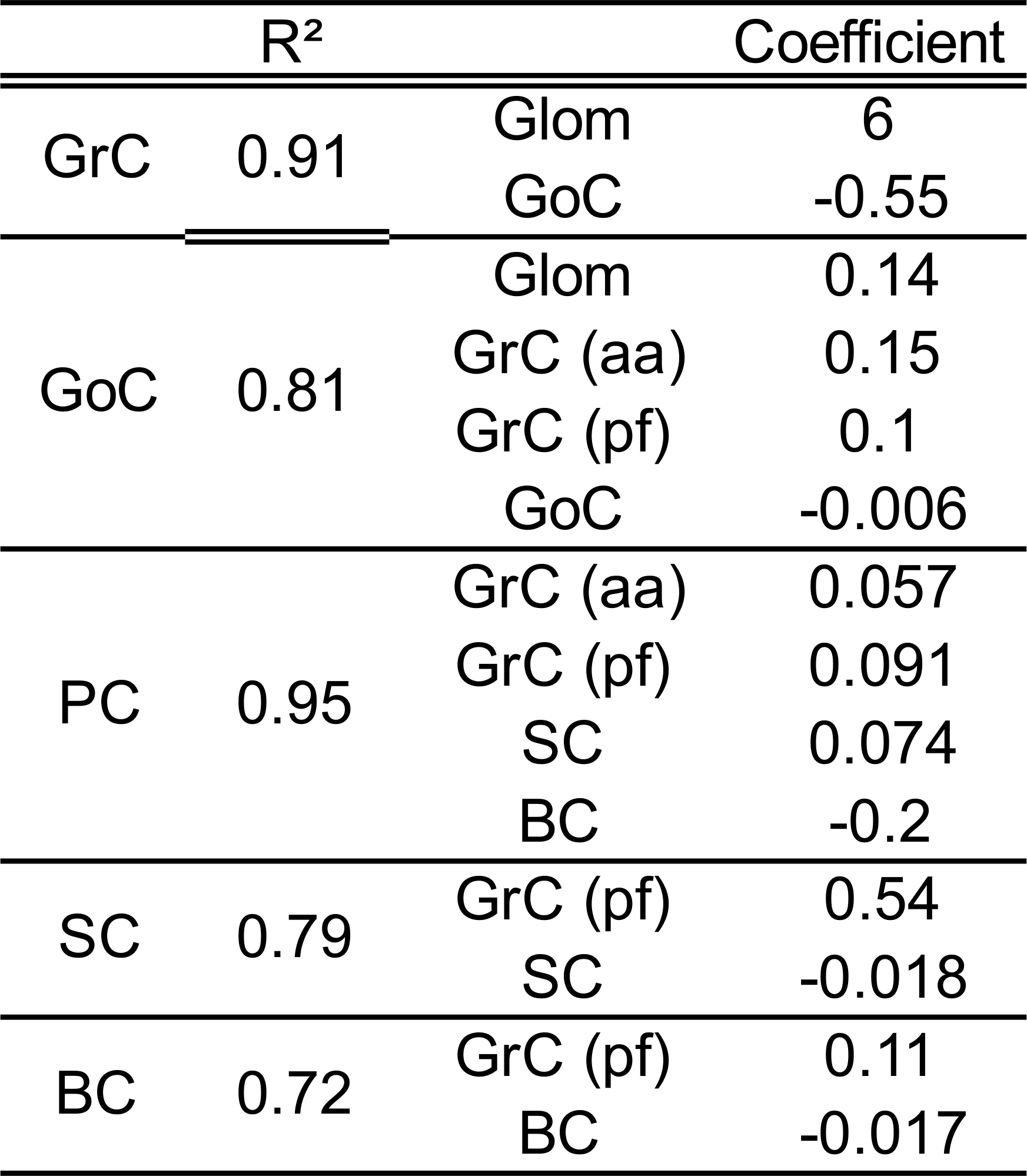
Input-output dependency of model neurons. Multiple linear regression between neuronal responses to the *mf* burst (firing rate during 40 ms after the stimulus onset) and the number of incoming spikes from the presynaptic neurons, averaged over 10 simulations. R^2^ and direction coefficients are reported.

**Figure S6.**
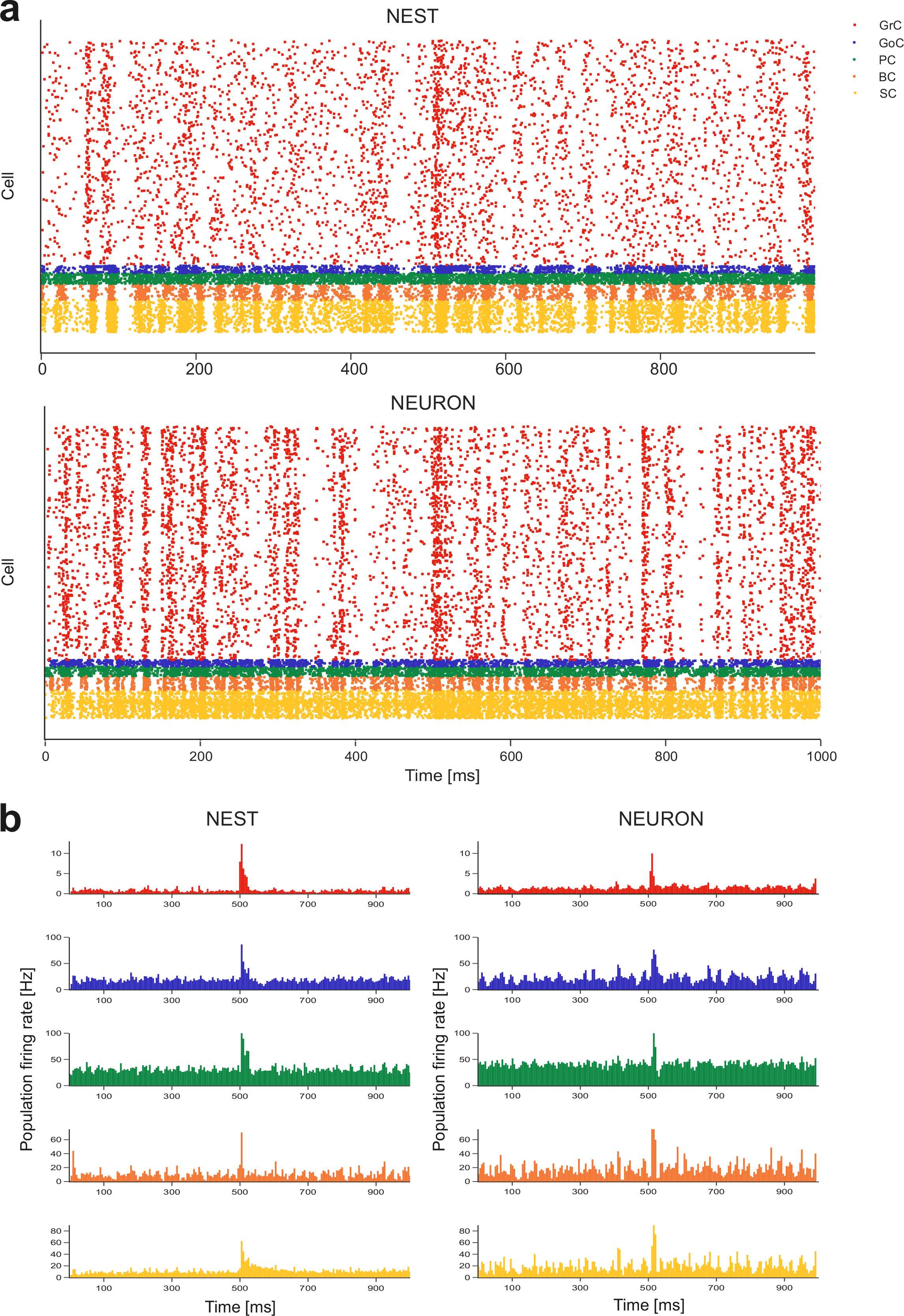
Comparison of NEURON to NEST simulators. The raster plots **(a)** and PSTH **(a)** that were generated using NEST and NEURON simulators are compared. In raster plots, just 10% GrCs are randomly sampled for clarity. The PSTHs show number of spikes/5 ms time-bins normalized by the number of cells, average of 10 simulations (as in Fig. 3b). The cerebellar network reconstructed using detailed neurons was simulated using mono compartment “single-point” neurons. The same stimulation protocol was applied (*mf* burst 20ms@200Hz over background noise; onset indicated by arrowhead). Network structure was unchanged, so that the number of synapses computed with detailed morphologies was maintained and applied to the soma of point neurons. All cellular mechanisms were taken from a previous NEST model including E-GLIF neuron models and alpha-based conductance synapses (Geminiani et al., 2018, 2019). So, what was lost was not just the electrotonic structure and the ionic channel distribution, but also the specific dendritic locations of synapses along with the dynamic mechanisms of neurotransmitter release. It should be noted from raster plots that single cell spike timing is visibly modified using NEST instead of NEURON, while the average burst response pattern visible in PSTHs remains similar for the various cell populations. This result highlights the impact of neuron microscopic properties and dendritic processing on single neuron computation and the maintenance of ensemble network dynamics in simplified network models.

**Movie S1. Impulsive response of the cerebellar network.** The movie shows the activation of a slab of the cerebellar network model by a train of 4 synchronized pulses @ 50 Hz to a bundle of 13 mfs (at time = 20 ms) riding over a background activity elicited by 4 Hz random stimulation to all *mf*s. Note the sparse activity in GrCs that suddenly aggregates into dense clusters activating the overlaying PCs and MLIs.

**Movie S2. GoC respons**e. The electrical activity of a GoC embedded into the cerebellar network (same cell as in Fig. 4c) is animated along with dynamic changes in molecular and synaptic variables. The movie shows 100 ms of simulation during random *mf* activity at 4 Hz with a burst delivered to 4 *mf* at 100 ms. Each variable is reported on the GoC morphology (using a colour scale) and shown at the bottom with traces referring to specific neuron compartments (using a moving time window). Three locations are shown for soma (teal), an apical dendrite (pink) and a basal dendrite (gold). (**Vm)** membrane potential traces in the soma and apical dendrite. **(Ca)** [Ca^2+^]_in_ in the basal and apical dendrite. **(AMPA)** synaptic current in the basal and apical dendrite (synapses from *mf*s, *aa*s, and *pf*s). **(NMDA)** synaptic current in the basal dendrite (synapses from *mf*s and *aa*s). **(GABA)** Synaptic current in the basal dendrite (synapses from GoCs).

